# Proton and alpha radiation-induced mutational profiles in human cells

**DOI:** 10.1101/2022.07.29.501997

**Authors:** Tiffany M. Delhomme, Maia Munteanu, Manuela Buonanno, Veljko Grilj, Josep Biayna, Fran Supek

**Affiliations:** Genome Data Science, Institute for Research in Biomedicine (IRB Barcelona), Barcelona, Spain; Radiological Research Accelerator Facility (RARAF), Columbia University, New York, USA; Catalan Institution for Research and Advanced Studies (ICREA), Barcelona, Spain

## Abstract

Ionizing radiation (IR) is known to be DNA damaging and mutagenic, however less is known about which mutational footprints result from exposures of human cells to different types of IR. We were interested in the mutagenic effects of particle radiation exposures on genomes of various human cell types, in order to gauge the genotoxic risks of galactic cosmic radiation, and of certain types of tumor radiotherapy. To this end, we exposed cultured cell lines from the human blood, breast and lung to intermittent proton and alpha particle (helium nuclei) beams at doses sufficient to considerably affect cell viability. Whole-genome sequencing revealed that mutation rates were not overall markedly increased upon proton and alpha exposures. However, there were changes in mutation spectra and distributions, such as the increases in clustered mutations and of certain types of indels and structural variants. The spectrum of mutagenic effects of particle beams may be cell-type and/or genetic background specific. Overall, the mutational effects of recurrent exposures to proton and alpha radiation on human cells in culture appear subtle, however further work is warranted to understand effects of long-term exposures on various human tissues.

## Introduction

Humans have a keen interest in travelling in space, from deep space exploration, to extraterrestrial colonization. In addition, the space environment was also considered for biomedical research purposes, for example the “Tumors in space” experiment aims to study the effects of microgravity and exposure to space radiation on tumor organoids, with implications to cancer risk and treatment^1^. Spaceflight, i.e. flight beyond the Earth’s atmosphere^2^, is usually defined as crossing the Kármán separation line i.e. 80-100 kilometres altitude above the sea level^2–4^. Above this boundary, the environment dramatically changes, notably in terms of microgravity, temperature and radiation^5^. Studies revealed the pathological consequences on human physiology from travelling in space and reported cardiovascular, musculoskeletal and immune changes^6^. However, a better understanding of space travel impact on the human body at the cellular and sub-cellular levels is needed, for instance how the genome integrity is affected. Important effects of space travel are exerted by the various types of radiation to which tissues are exposed outside the Earth’s atmosphere and magnetosphere. Those include the solar radiation, the radioactive environment of the planet, and galactic cosmic radiation (or rays; GCR).

GCR originates mainly from the Sun or outside of the Solar system but within the Milky Way galaxy (there exist also the extra-galactic cosmic radiations, which are less relevant as a space travel hazard due to their low flux and extremely high energies). The GCR encompasses photon-based radiation as well as particle radiations, including neutrons and charged particles; among the latter, protons and helium nuclei are the most abundant. Cosmic radiation has emerged as an issue of concern, since exposure to radiation may increase cancer risk in astronauts^7, 8^. A better understanding of the mutagenic mechanisms of various cosmic radiation types upon human DNA would be helpful to understand the basis of the increased risk of cancer or of reproductive harm resulting from space travel.

Photons are the better-studied component of the GCR, since they consist of gamma rays and X-rays, whose biological impact was studied in other contexts apart from GCR. For instance, gamma rays were released during the Chernobyl nuclear power plant disaster, prompting studies of how this type of radiation impacts human genome stability. The genomic profiles of thyroid carcinomas in irradiated children post-Chernobyl contained similar driver mutations and gene fusions as in non-radiation-associated thyroid tumors^9^. However, there was a radiation dose-dependent increase in the fusion driver events, as well as a general radiation-associated increase in certain types of structural variants (rearrangements)^10^. Consistent with a mutational impact that can generate somatic driver mutations, ionizing radiation-generating incidents (e.g. the Chernobyl disaster, and also the Hiroshima and Nagasaki A-bomb attacks) also increased the risk of developing cancer, proportionally to the dose of exposure^11, 12^.

In addition to nuclear incidents, radiation exposures can result from medical uses. Commonly this involves X-rays, used for both diagnostic purposes, and also in applications to tumor radiotherapy. The impact of therapeutic X-ray exposure on human DNA was examined via genomic analyses of radiation-associated second malignancies and of post-treatment metastatic tumors, reporting an increase of small and large deletions, and also certain mutational signatures^13, 14^. For instance, this entails a particular signature of indels (the PCAWG Signature ID8) characterized by larger deletions (≥ 5-bp) without flanking micro-homology; this was previously linked to double-strand break repair^15^. Consistently with their mutagenic effect, medical use of X-rays has been reported to increase cancer risk^16^.

Regardless of the source of the radiation – either the cosmos, nuclear accidents or medical uses – it is not known if ionizing radiation consisting of charged-particle radiations, such as protons or alpha rays, would have similar mutagenic effects to more commonly studied photons (X-rays and gamma-rays). Generally, all types of ionizing radiations, including gamma-rays, X-rays, energetic charged particles (protons, alpha particles [helium nuclei] and heavier ions) and neutrons have been classified as human carcinogens by the World Health Organization (WHO)^17^. This suggests mutagenic impacts of various radiation types, however research on the specific mutational footprints of charged particles on the integrity of the human genome has been limited.

These are of interest because during transit beyond low Earth orbit, every cell nucleus within an astronaut’s body is traversed, on average, by an energetic proton every few days, and by a helium nucleus once every few weeks. Mutational effect of proton radiation is of interest also for reasons other than space travel. This type of radiation is increasingly adopted in the clinic to treat cancer^18^ , and appears promising for toxicity profiles^19–22^. Proton therapy may reduce the health risks (compared to X-rays) of secondary malignancies^23^, and of severe radiation-induced lymphopenia^24, 25^. The above provides motivation for studying the mutagenic effects of protons, and of helium nuclei (alpha) radiation exposures, which have the potential to result in long term health effects in exposed individuals.

The activity of mutational processes in DNA, which may arise from both endogenous factors such as DNA repair deficiencies and exogenous factors such as tobacco smoking, can be captured by mutational signatures^26^, mathematical constructs that describe the differential frequencies of mutation types across many genomes. Such signatures are commonly based on trinucleotide spectra of single nucleotide variants (SNVs), however they can also be based on indels, structural variation (rearrangements), or mutation clusters^15, 27, 28^. SNV and indel mutational signatures in human cancer are well characterized and organized in comprehensive catalogues^15, 29^, including also signatures in tumors pre-treated with ionizing radiation^13, 14, 30^ (here referring to photons).

In addition, some genomic studies have reported mutational signatures on experimental models, including experimentally-induced mutational signatures of (non-ionizing) UV radiation in human cell lines^31^, and also in cultured cells from healthy human tissues^32, 33^. Additionally, signatures of ionizing radiations were reported in exposed mice^34, 35^ and in the worm *C. elegans*^36^; both reported SNV signatures enriched in C>T transitions^15^.

In order to increase our understanding of the potential impact of particle radiation encountered during space travel on human tissues, it is important to systematically, in a genome-wide manner, examine the DNA mutations arising from the exposure in different human tissues. To this end, we analyse genomes three human cancer cell lines, A549 (lung adenocarcinoma, epithelial cells), HAP1 (chronic myelogenous leukaemia, blood cells) and MCF7 (breast carcinoma, epithelial cells). Those cell lines were irradiated with two types of particle radiations – proton and helium fluxes – in order to identify the mutational footprints induced and their potential cell type-specificity.

## Results

### Determining proton and helium ion fraction, and irradiating the cell lines

The dosage of radiation was determined experimentally in order to achieve between 40% and 50% lethality (corresponding to 50%-60% clonogenic survival), independently across the two types of cosmic rays (Figure 1A-B).

**Figure 1.**
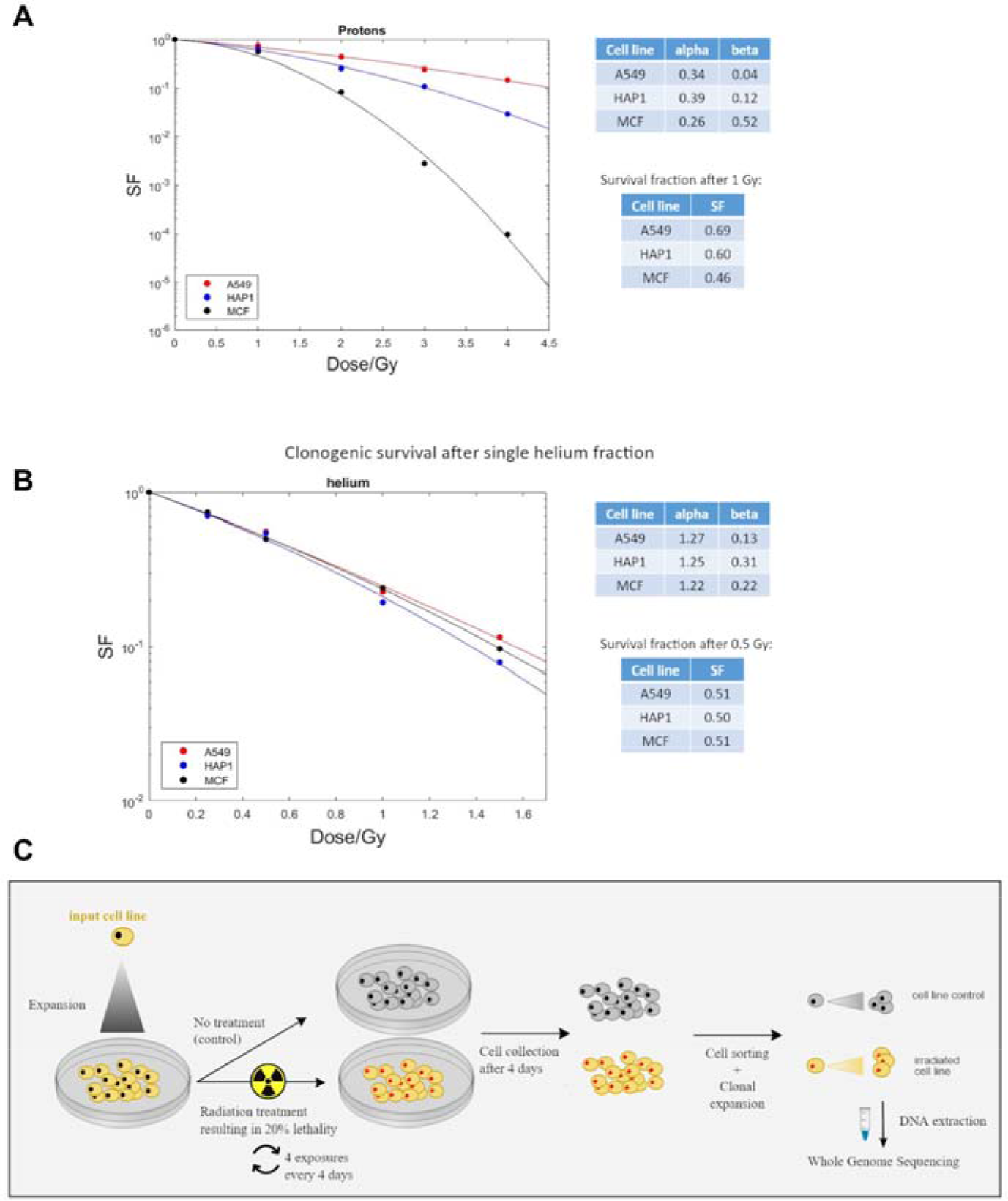
Overview of the experimental design. **A-B.** Experimental identification of radiation dose in order to achieve ∼50% of cell lethality (i.e. ∼50% of cell survival) independently across the three cell lines and across the two types of radiations. SF, survival fraction. C: Schematic overview of the experimental design of the study. A549, MCF7 and HAP-1 cell lines were exposed to GCR (protons or helium ions). GCR: Galactic Cosmic Radiation. Single cells from treated and untreated conditions were separated by FACS and clonally expanded. DNA from single-cell derived populations was extracted and subjected to whole genome sequencing.

This led to a dose of 0.5 Gy for helium fluxes and a dose of 1 Gy for proton fluxes. After an expansion of the three cell lines collected (A549, HAP1 and MCF7), the cells were irradiated using the 5.5 MV Singletron accelerator at the Radiological Research Accelerator Facility (RARAF; see Methods for details about the procedure, with the corresponding dosage). We exposed each cell line four times every 4 days to allow cell recovery between each exposure to facilitate mutation accumulation.

After the final latency period of 4 days, the cells were collected, sorted in a 96-well plate, and a clonal expansion was performed to obtain colonies with a near-identical genome to facilitate identifying mutations (Figure 1C). Finally, the DNA was extracted from those colonies and whole-genome sequenced (WGS) on an Illumina NovaSeq 6000 machine. Details of the following bioinformatics analysis are described in Methods.

### Overall burden of various mutation types generated by cosmic radiations

After the GCR treatments, the cell line genomes contained newly-acquired point mutations (single nucleotide variants, SNVs), short insertions and deletions (indels), and structural variants (SVs) in all three cell lines assayed, HAP1, A549 and MCF7, and regardless of the radiation type, protons or helium ions. As a control, we considered clones generated from sham-irradiated (i.e. untreated) cells for all three cell lines; these clones also accumulated certain numbers of SNVs, small indels and SVs, consistent with them undergoing cell divisions and also potentially being exposed to various stresses during handling.

The global number of observed mutation events (SNVs and indels) does not markedly differ between conditions (Figure 2A), suggesting that proton and alpha radiation exposure is not grossly mutagenic to cultured human cells using the fractionated dosing regimen applied herein. It bears mentioning that we detected higher amounts of SNVs in the MCF7 cell line (roughly double) than in the two other cell lines after exposure to either particle as well as in the MCF7 untreated cells (Figure 2A).

**Figure 2.**
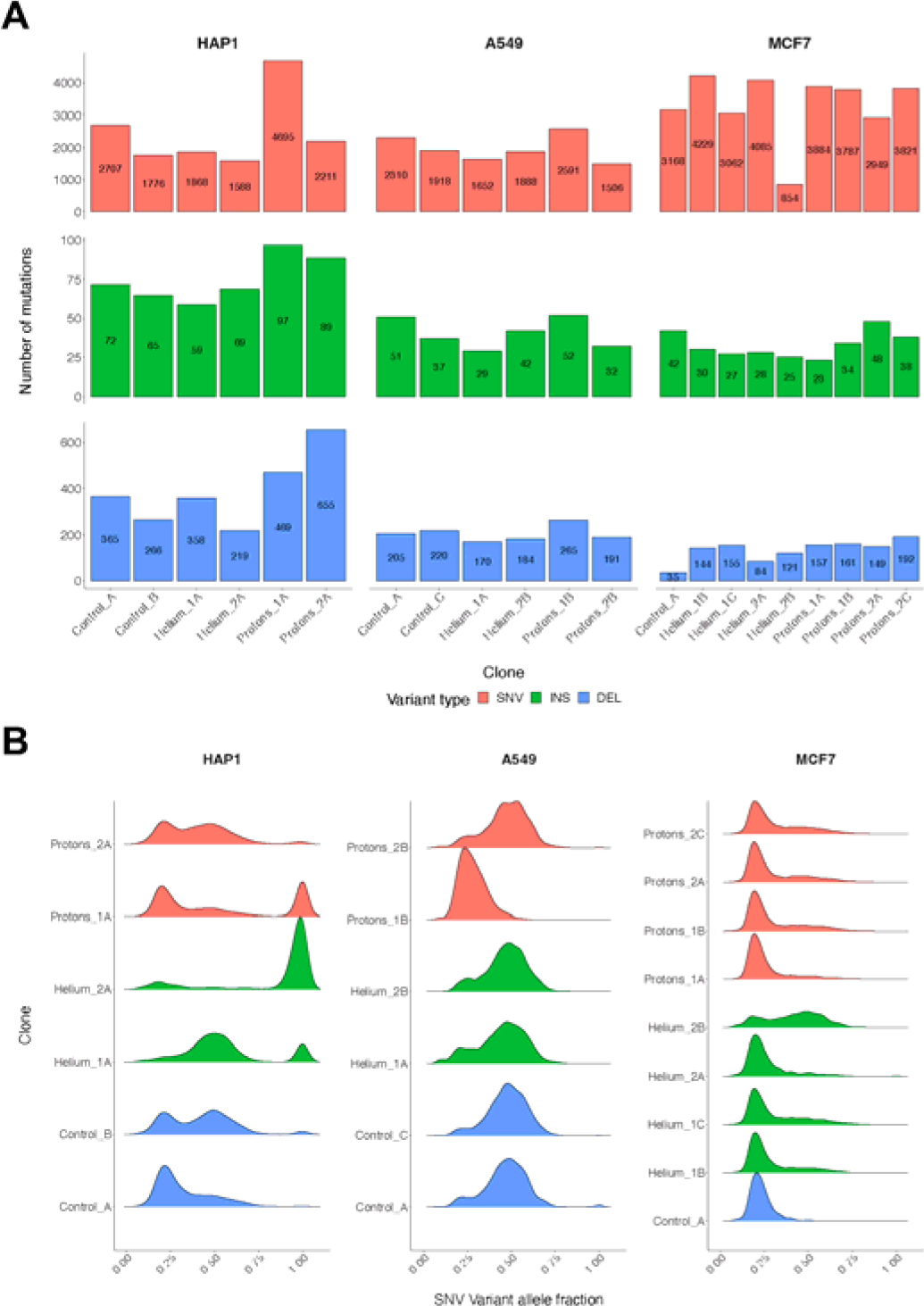
General description of point mutations detected in the clones. **A.** Distribution of the number of SNVs, small insertions and small deletions per clone in each of the three analyzed cell lines. **B.** Distribution of the variant allele fraction (VAF) of detected point mutations for each clone of each cell line. A VAF peak around 50% or 100% would correspond to a selection of precisely one cell from the cell culture, whereas a peak close to 25% would correspond to the isolation of two cells or cell line polyploidy.

This may indicate a mutator phenotype in our MCF7 cell line, possibly due to a DNA mismatch repair (MMR) failure (see mutation signature analysis below). However this is difficult to ascertain since the absolute number of mutations between cell lines/ conditions cannot be compared precisely, due to a potentially variable number of cell cycles the cells have undergone during the experiment. We further noted an increased number of indels, particularly deletions, in the HAP1 cell line compared to other two cell lines (Figure 2A). Because this was observed also in the control cells again this suggests that our HAP1 cells have an intrinsically higher rate of accumulating indels, often 1nt deletions at homopolymers, suggesting a type of microsatellite instability.

To comprehensively assess the differences in mutation burden between different conditions, we implemented a randomization test (see Methods for details, Supp. Figure 1). In all irradiated cell lines, specifically the proton treatment generated the clone with the highest number of indels in its genome, compared to the other conditions (Figure 2A). This trend is also visible in the randomization results, where the proton-treated clones have more indels compared to helium and untreated groups (unadjusted p=0.078, p=0.273), especially clear (though not statistically significant after FDR correction) in the HAP1 cell line (unadjusted p=0.047, p=0.031).For 2 out of 3 cell lines, the proton treatment also generated the clone with the highest number of SNVs and generally appeared to generate more SNVs across all cell lines compared to the other treatment groups (p=0.096 for control, p=0.074 for the helium comparison). The indel-enrichment trend for proton-treated clones was more prominent for deletions than for insertions (Figure 2A, Supp. Figure 1, panel “IDs_del_ins_ratio” p=0.055). An enrichment of deletions, as we observed in the proton-treated clones, has recently been highlighted in ionizing radiation-associated tumors^13, 14^ which are treated with photon radiation, typically X-rays. The globally most-mutated clone is a single proton-irradiated sample of the HAP1 cell line, which contains 4695 SNVs (Figure 2A), compared to 2707 and 1776 in the control (non-irradiated) HAP1 samples. Proton flux also generated the highest number of insertions in the HAP1 cell line (97 and 89 detected in the two proton-treated replicates, compared to 72 in the control), and also deletions (469 and 655 detected in proton-treated, compared to 365 in the controls). For this type of alteration, there was a more modest impact of the proton treatment in the A549 and MCF7 cell lines (Figure 2A). We also note considerable variation in the mutation burden of proton-treated samples between clones (Figure 2A). Overall, this suggests that mutagenic impact of proton irradiation on SNVs, insertions and deletions is evident however it also may be rather variable depending on cell-type and stochastic factors that differ between individual cells.

Variant allele fraction (VAF), the proportion of sequencing reads that contain a particular variant, is a proxy for the proportion of cells in the sequenced population that harbor the mutation. In our experiments, we aimed to select a single treated cell and expand it into a clone, thus it is expected that the VAFs observed should be centered at 0.5 for heterozygous diploid genome segments). In practice, three different types of VAF patterns were observed (Figure 2B). One peak at 0.25, which suggests that either two diploid cells were collected at the time of bottlenecking or alternatively, that the genome or its segment may be tetraploid; we observed this commonly for the MCF7 cell line, which is indeed known to be hypertriploid to hypotetraploid^37^. Second pattern is one peak at 0.5 which suggests that one cell was collected and that mutations were in a diploid segment; this was common for the A549 cells. Finally, a third pattern, a peak at 1, was observed for the HAP1 cell line, as expected due to the unusual genome-wide haploid state of that cell line (Figure 2B); this cell line can grow as haploid or diploid and may switch spontaneously^38^ and indeed we observed considerable number of VAFs <=0.5 in HAP1 genomes (Figure 2B). The VAF distributions in some HAP1 clones e.g. PR1A suggest that a whole-genome doubling may have occurred. We note that one helium-treated MCF7 clone exhibited an unusual VAF distribution, as well as outlying (low) SNV burdens (Figure 2A), suggesting technical artifacts in this experimental replicate and we thus excluded the WGS data from this MCF7 sample from further display, whilst still retaining it in the main analyses.

### Identification of SNV trinucleotide mutational signatures

Upon classifying SNVs into 6 different categories, tallying them DNA strand-symmetrically, i.e. C>A, C>G, C>T, T>A, T>C, T>G, the most prevalent class of SNVs in all cell lines/clones was C>A (Figure 3A). One important cause of C>A transversion mutations in cancer genomes is oxidative damage to guanine (Signatures SBS18 and SBS36 as reported^26^). Moreover, an abundance of C>A mutations was observed in recent mutation-accumulation experiments on human cell lines similar to ours^31, 39^, plausibly due to exposure to atmospheric oxygen during cell culture conditions. Control (unirradiated) clones also had high C>A exposures.

**Figure 3.**
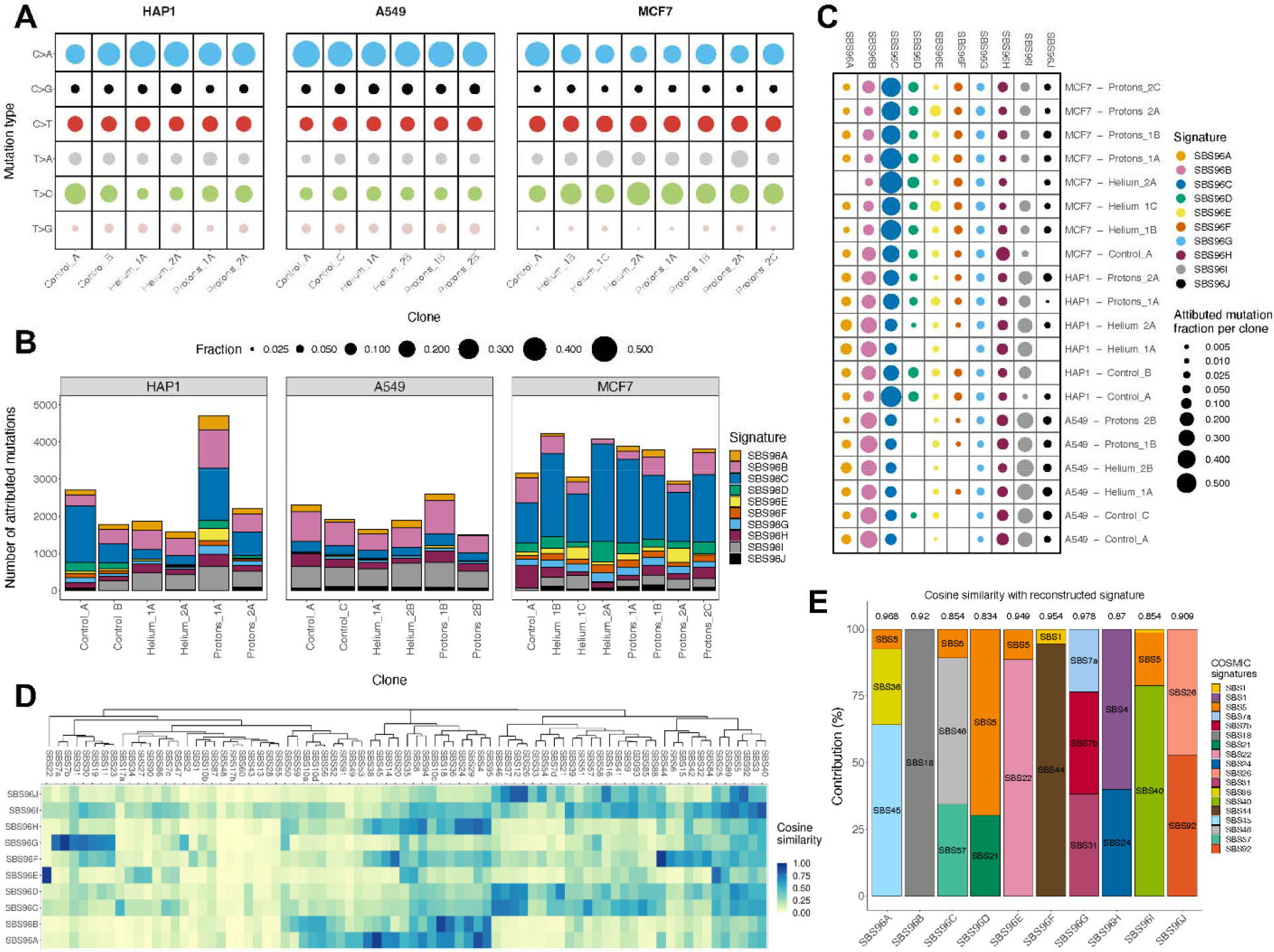
Mutational signatures of SNVs in irradiated human cell lines. **A.** Break-down of point mutations into the 6 main types of mutations (*i.e.* C>A, C>G, C>T, T>A, T>C, T>G), relative contributions per clone in each cell line. **B.** Number of point mutations attributed to each of the 10 different extracted SNV mutational signatures, for each sample in our cohort). **C.** Exposures of all extracted SNV mutational signatures expressed as a fraction of the total mutation burden in each clone. **D.** Cosine similarities between PCAWG mutational signatures and our 10 extracted SNV signatures. The mutational signature dendrogram was generated with the hclust function from the stats R package, with the agglomeration method set to “complete”. E. Decomposition of extracted signatures into PCAWG signature spectra. Values at the top of each bar represent the cosine similarity between the original signature profile and the reconstructed spectrum.

Additionally, we also observed considerable numbers of the C>T and T>C transition mutation types (Figure 3A), which is consistent with commonly observed mutational signatures in dividing cells, namely the ‘clock-like’ mutational signatures SBS1 and SBS5/SBS40^15^. These transition mutations were particularly abundant in the MCF7 genomes, suggesting a mutational process specific to that cell line.

To further investigate mutational processes, the SNV mutations were classified into 96 categories, taking into account both the observed class of mutation and the trinucleotide context (A_A, A_C, …, T_T), in order to infer the 96-component mutation spectrum of SNVs for each cell line and each clone (Supp. Figure 10, Supp. Table 1).

We extracted *de* novo mutational signatures from our irradiated and control clone WGS via the SigProfiler tool^15^. To identify similarities with known mutational processes from previous experiments on human cell line models, our data was pooled with previous genomic data: (i) 155 cell line genomes treated with various mutagens, largely chemicals, in which a specific SNV signature was able to be identified as well as gamma-irradiated genomes (with no identifiable signature)^31^, and (ii) an additional set of 38 cell line genomes generated from CRISPR-Cas9 gene knockouts of 9 DNA repair genes reported to exhibit mutagenesis upon knockout^39^. By a joint analysis of this mutation accumulation data, we identified 10 independent mutational signatures (see Methods) based on SNVs (here called SBS [single base substitution]) (Supp. Figure 2A) and examined their exposure across all clones in our cohort (Figure 3B-C. Supp. Figure 2B-C),

In order to attribute a potential aetiology to each signature, we computed the similarity of their spectra with the catalog of known somatic signatures, here referred to as PCAWG signatures^15^ (Figure 3D; of note, these signatures are often referred to in the literature as “COSMIC”, by the name of the database, however this is unrelated with “cosmic radiation” and this previous catalog does not contain known signatures of cosmic radiation). Additionally we used a “decomposition” approach to model our signatures as mixtures of known PCAWG signatures (Figure 3E). Finally, we studied aetiologies of our 10 signatures by considering the activities of these signatures in particular samples with known DNA repair deficiency or with exposures to known mutagenic chemicals or radiation from two previous studies^31, 39^. (Supp. Figure 3A)

### Mutational signatures observed across our and prior cell line experiments

The most abundant signatures in the pooled dataset were SBS96A and SBS96B. The SBS96A signature was assigned to (Figure 3D-E) both PCAWG SBS36 (a C>A signature of oxidative damage to DNA, associated with failures in base excision repair) and to SBS45 (C>A-rich signature, likely an experimental artifact due to 8-oxoG generated *in vitro* during preparation of DNA for sequencing^40^); the signature activities were high in the experimental exposure to certain PAHs (polycyclic aromatic hydrocarbons)^31^ (Supp. Figure 3A). The SBS96B was assigned to the PCAWG signature SBS18, a signature resulting from damage to guanines by reactive oxygen species^15, 41, 42^. This signature was highly active in a *OGG1* knockout cell line genome (Supp. Figure 3A), consistent with the known function of base excision repair protein OGG1 in mending nucleobase oxidative damage^43^ and additionally in treatments with e.g. potassium bromate (an oxidizing agent) and gamma radiation (Supp. Figure 3A).

Next, SBS96C, a signature abundant in the MCF7 cell line genomes in our WGS dataset (Figure 3B), was not assigned to a single reported signature in PCAWG catalog (Figure 3D), but instead the decomposition models it, at modest accuracy, as a mix of SBS46 (T>C transitions in various contexts, with some C>T transitions) and SBS57 (T>C and T>G mutations in TTT) (Figure 3E). The SBS46 and SBS57 are noted to be possible sequencing artifacts in the COSMIC database. However, we infer (see below) that this is a *bona fide* mutational process not previously reported; a defect in one or more of the DNA repair pathways might generate the signature SBS96C in copious amounts.

Signature SBS96D was also likely to be not seen in previous PCAWG signatures (it is modeled, at low accuracy, as a mix of the ubiquitous, unknown aetiology SBS5, and SBS21, one signature of defective DNA mismatch repair (MMR). This signature dominated by T>C and C>T transitions, had high exposures in previous samples treated with methylating agents (ENU, MNU, TMZ, 1,2-DMH; Supp. Figure 3A), suggesting the SBS96D originates from alkylating DNA damage.

The following four signatures, SBS96E–H, appear to have clear aetiologies. SBS96E corresponded closely to the PCAWG signature SBS22 (Fig 3D-E), resulting from an exposure to aristolochic acid (Supp. Figure 3A), an agent generating bulky nucleotide adducts, resulting in T>A transversions. SBS96F was assigned to the PCAWG signature SBS44 and to the *MLH1*, *MSH2* and *MSH6* gene knockouts^39^, resembling a signature of a defective DNA mismatch repair. SBS96G was assigned to the known mutational signatures associated with ultraviolet light exposure (SBS7a, SBS7b), and had high exposure in the simulated solar (i.e. non-ionizing, UV-containing) radiation treatment of a cell line^31^ (Supplementary Figure S3A). SBS96H is a mix of SBS4 (tobacco smoking-associated) and SBS24 (aflatoxin-associated), and was seen in cells exposed to BPDE and PhIP chemicals in previous data, thus reflecting mutagenesis due to bulky adducts in DNA. These signatures, SBS96E–H, tend to generate fewer mutations in our samples; the latter, signature H, is somewhat more abundant in both treated and untreated samples (Figure 3B-C).

Signature SBS96I likely corresponds to the ubiquitous, clock-like signatures of unknown origin SBS40 and SBS5 (although we note inaccurate spectrum reconstruction, Figure 3E) and is abundantly present in our samples. In previous data, we note a link to the knockouts of the ubiquitin ligase *RNF168* and exonuclease *EXO1* (Supp. Figure 3A), involved in DNA damage signalling and repair respectively. Signature SBS96J contains SBS26 (DNA MMR deficiency) in its spectrum deconstruction (Figure 3E), and it is seen in the knockouts for the *PMS2* gene of the DNA MMR pathway (Supp. Figure 3A); however it also contains SBS92 (tobacco smoke metabolites exposure) and is found in genomes treated with 6-nitrochrysene; thus its underlying mechanism is unclear or it may represent a mixture of mechanisms.

### Abundant SNV mutational signatures vary in activity across cell lines

We further considered signature activity in a particular sample in relation to the treatment and/or to the cell line of origin. In order to systematically compare between these groups, we implemented a randomisation strategy (Supp. Figure 4A, see Methods for details).

Firstly, we consider the three most abundant signatures in our cell line data: SBS96B (oxidative DNA damage, ∼SBS18), C (possible DNA repair defect and/or artifacts), and I (background mutagenesis, ∼SBS5/40). These were found at various levels in the three cell lines and in the majority of the samples of each line and were also seen in the non-irradiated control samples (Figure 3C).

The canonical signature of reactive oxygen damage, SBS96B, was detected also in the non-irradiated clones, probably resulting from DNA oxidation reactions during cell culture. SBS96B, however, does trend towards a higher activity (Supp. Figure 4A; p=0.079; n.s. upon FDR adjustment) in the proton irradiated *versus* helium irradiated cell lines (Figure 3C; Supp. Figure 4A). This trend is seen when considering the cell lines jointly, and also considered individually in comparisons of proton *versus* helium treatments (Supp. Figure 4A). Our data is consistent with a mechanism of more rapid formation of reactive oxygen species as a consequence of proton radiation, however it could also result from other factors (a higher rate of DNA damage resulting from the reactive oxygen, and/or compromised repair of oxidized DNA upon proton radiation). Irrespective of the treatment, this signature was seen at different abundance across the cell lines, in order A549>HAP1>MCF7.

Our SBS96C signature, modeled as a mixture of two known PCAWG artifact signatures, is however unlikely to be an artifact since it is present differentially in all MCF7 cell line WGS *versus* the other two cell lines – A549 and HAP1 – where it is rare (Supp. Figure 4A, p=0.00008 and p=0.185, respectively). It does not associate with radiation exposures. Since all our samples and resulting data were treated equally, it does not seem likely that a sequencing artifact would arise in data from one cell line, while largely absent in the other two. Additionally, the attempt of reconstruction of SBS96C as a mixture of the known artifact signatures SBS46 and SBS57 was not highly accurate (Figure 3E, cosine similarity = 0.854), further supporting the idea that this is a genuine mutagenic process, possibly an acquired DNA repair deficiency, SBS96D was found in abundance in all samples of the MCF7 cell line including the control, and interestingly also in two clones from the HAP1 cell line including one control. We infer this may be a form of MMR deficiency, based on the co-occurrence with indel signatures (see below), possibly combined with other DNA repair deficiencies thus explaining the unique SBS spectrum not observed in previous cell line data considered here.

Another abundant signature was SBS96I, referring to SBS40-like background mutagenesis, which was not associated with particle radiation treatments in the 3 cell lines considered jointly, however its abundance was strongly variable across cell lines in the order of A549>HAP1>MCF7 (Figure 3B-C, Supp. Figure 4A), consistently as SBS96B above, potentially indicating a clock-like nature of signatures SBS96I and SBS96B in our experimental setup, inferred from a slow division rate of MCF7 cells (not shown).

### Signatures not associated with the particle radiation treatments

SBS96F, the DNA repair deficiency-associated signature that strongly associated with deletion of the MMR genes *MLH1*, *MSH2* and *MSH6* (Supp. Figure 3A) was not highly active in our samples; instead, it was seen mostly in the external datasets we integrated. We do note some enrichment of SBS96F in the MCF7 cell line, however no association with either type of radiation treatment was observed in any of the 3 cell lines (Supp. Figure 4A). Thus, particle radiation exposure does not commonly select for cells with a deficient MMR system, as is the case for some chemical exposures that generate mismatch-like DNA lesions^44^. Consistently, SBS96J, which might be associated with DNA repair inefficiency via *PMS2* deletion (Supp. Figure 3A), is overall present at low levels but more in A549 and MCF7 than in HAP1 (Figure 3B-C), and appears not associated with treatment in 2 cell lines (some association noted in MCF7 cells only; Supp. Figure 4A). Overall, signatures of deficient MMR in our data appear unrelated with alpha or proton radiation exposure, however they do differ between cell lines.

The SBS96D signature, likely originating from methylating DNA damage, is not associated with particle radiation treatment, but differs considerably between cell lines in activity (MCF7>HAP1>A549; Supp. Figure 4A). Similarly, the SBS96A signature, likely resulting from exposure to certain types of oxidative DNA damage and/or failures to repair the damage, does not show association to our particle radiation treatment overall nor in 2 of the cell lines (there may be some signal in HAP1 cells; Supp.Figure 4A).

The mutational signature of the non-ionizing UV radiation SBS96G, was, expectedly, detected at only minor levels (≤ ∼5% of observed mutations) in our data (Figure 3B-C). While overall it was slightly (albeit positively) associated with proton radiation treatment (Supp. Figure 4A, p=0.075 for control vs protons; n.s. after FDR adjustment), this association was seen only in one cell line..

### Signatures with tentative links to particle radiation treatments

In addition to the common C>A signature of oxidative stress SBS96B mentioned above, we noted some evidence for radiation treatment association in our cell line experiments with two other signatures. Firstly, the T>A transversion signature SBS96E (Figure 3B-C), associated with bulky DNA adduct-forming mutagen exposures, while present overall at low levels in our data (somewhat higher in MCF7 cells) was modestly positively associated with proton treatment (Supp. Figure 4A, p=0.097 for control vs protons). Similarly, another signature associated with bulky DNA adducts, the SBS96H (seen in previous BPDE-treated samples and similar to tobacco-smoking SBS4), was consistently trending towards positive association with proton treatment compared to helium treatment (Supp. Figure 4A, p=0.160 for helium vs protons; n.s. after FDR adjustment). For the SBS96E and SBS96H associations with particle radiation, these trends were seen in at least 2 out of 3 cell lines and are thus plausible, however it should be noted that the effects were modest and did vary across the cell lines. One possible explanation is that there is genuine variation between cell types and/or genetic background in the mutational signatures of particle radiation.However it is also possible that the subtle effect of particle radiation on SNV spectra is in fact universal but did not cross a detection threshold in one of the cell lines.

The mechanisms that underlie signatures SBS96B, E and H have a common thread: these mutational signatures are thought to result from large, replication-blocking DNA lesions. Thus, proton/helium radiation effect on DNA can mimic the mutational footprint of mutagens generating bulky adducts. Speculatively, such a DNA lesion might be generated by the radiation causing intrastrand nucleotide cross-links, or by cross-linking DNA with proteins. Since these SNV mutational signatures were found in both protons and alpha rays treatment groups (of note, more so in the former), it does appear broadly relevant to various particle radiation types present in GCR. Moreover, it is expected to result from other sources of particle radiation (e.g. in cancer radiotherapy using protons) in the susceptible cell types and/or genetic backgrounds.

Based on repeated NMF runs on subsampled WGS data, we demonstrate that including a large amount of data from two prior studies^31, 39^ in our signature extraction dataset, as described above, does not have undue influence on the NMF point mutation signatures identified here (see Methods for description; Supp. Figure 5).

### Mutational signatures of indels across cell types and conditions

Small insertions and deletions (indels) can be classified into 83 types (the ID83 categorization, based on PCAWG indel signatures^15^). In brief, the indels are separated into 1-bp or longer indels, insertions or deletions, at DNA repeats or not, and finally deletions with micro-homology are counted separately; these classes can further be divided into subclasses based on the gained/lost nucleotide, or based on the alteration length. Each observed indel is assigned to one category, thus providing the mutational spectrum of indels for each cell line and each condition (Supp. Figure 11, Supp. Table 1).

In order to estimate the potential indel signatures as footprints of the helium and alpha radiation treatments, we ran the SigProfilerExtractorR tool on indel count matrices. The matrices were derived from the same pooled dataset, consisting, as for SNVs discussed above, of the genomes from our irradiated MCF7, A549 and HAP1 cell lines, as well as genomes of mutagen-treated and DNA repair deficient cell lines from previous studies^31, 39^ (see Methods for details).

Four indel signatures were identified: ID83A, ID83B, ID83C and ID83D (Figure 4A), as well as the activity (“exposure”) of each in our experimental cell line samples (Figure 4B). We also compared the number of small insertions and deletions in each treatment group, for each cell line (Figure 4C). To infer potential underlying mechanisms, we also examined the estimates of the signatures’ activity in previous DNA repair knockouts and the mutagen-treated samples^31, 39^ from which they were extracted (Supp. Figure 3B), as for the SNV signatures above. ID83A, consisting of 1nt deletions at A:T homopolymers closely matched the PCAWG ID2 signature (Figure 4D-E) and was abundant in DNA mismatch repair deficiency (*MSH2*, *MLH1* and *MSH6* knock-outs) (Supp. Figure 3B). Its sister signature isID83C , consisting mainly of 1nt insertions at A:T homopolymers, matching the PCAWG ID1 (Figure 4D-E), suggested to be related to slippage during DNA replication^15^. While ID83A contributes highly to the mutation burden of our samples, ID83C is responsible for a small number of mutations in HAP1 samples. Regarding the association to treatment, while the ID83A and C did display a weak positive association for protons, this was mainly seen in a single cell line (HAP1 in both cases).

**Figure 4.**
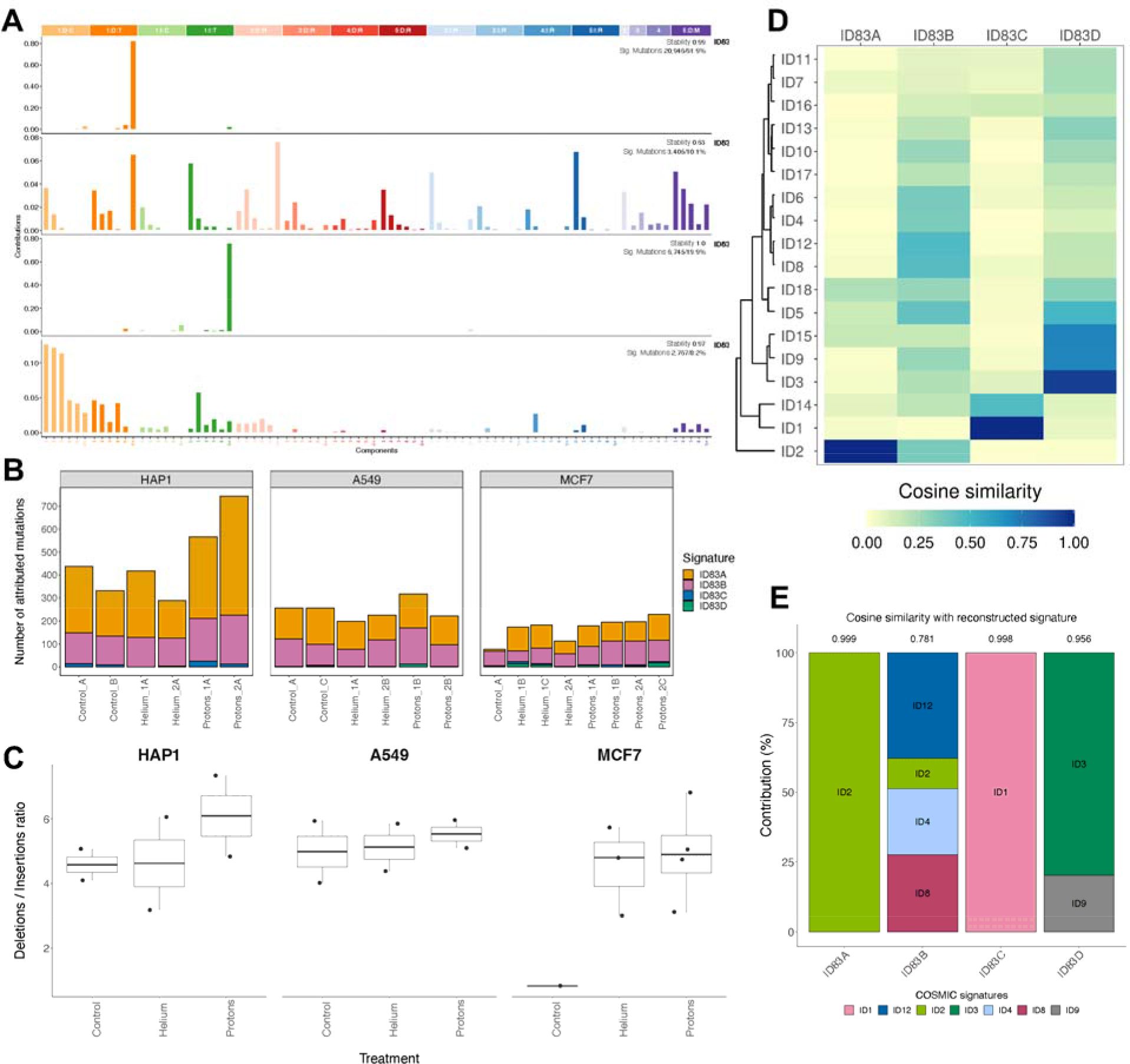
Mutational signatures of small insertions and deletions in irradiated human cell lines. **A.** Mutational profiles of the 4 indel signatures extracted using the SigProfiler ExtractorR algorithm. Indels were classified into 83 distinct categories following the procedure described in Alexandrov *et al.*^15^. **B.** The number of indels attributed to each of the 4 different signatures, for each sample in our cohort. **C.** Ratio of all small deletions over all small insertions, pooled by treatment for each cell line. D. Cosine similarities between PCAWG indel mutational signatures and our 4 extracted ID signatures. The mutational signature dendrogram was generated with the hclust function from the stats R package, with the agglomeration method set to “complete”. E. Decomposition of extracted signatures into PCAWG indel signature spectra. Values at the top of each bar represent the cosine similarity between the original signature profile and the reconstructed spectrum.

The remaining 2 signatures, ID83B and ID83D, were more interesting. ID83B was modeled as a mixture of 4 PCAWG signatures (Figure 4E), with higher weights of the ID12 (unknown aetiology) and ID8 (ionizing-radiation associated signature)^30, 45^ and ID4 (unknown aetiology). The ID83B spectrum consists of various lengths (1 bp to 5 bp with similar frequency) insertions and deletions outside of DNA repeats, as well as a notable component of deletions with short flanking microhomology (mostly 1-2 nt; we note this contrasts with the microhomology deletions resulting from homologous recombination deficiencies, which tend to be 2+ nt^46^). The activities of ID83B in prior cell line WGS data correspond to exposures to UV radiation and to propylene oxide and furan (Supp. Figure 3B). In our experiments, the ID83B signature was overall abundant and modestly enriched in proton-treated samples when considering the 3 pooled cell lines (Supp. Figure 4B, compared to helium-treated - p=0.017; there was a weak trend when compared to untreated - p=0.18); consistent direction of effect was observed in all 3 individual cell lines (Supp. Figure 4B).

Secondly, the ID83D signature consisted mostly of short 1-bp and 2-bp deletions outside of homopolymeric repeats and an additional minor component of ≥5bp deletions with short flanking micro-homology. This was found to be active in a cell line with *RNF168* knockout ^39^, a gene encoding a DNA damage signalling protein (Supp. Figure 3B), in cells treated with PAHs from tobacco smoke, (BPDE, DBADE and B[a]P, Supp. Figure 3B) and consistently it contained a component of PCAWG ID3 (Figure 4E), found in tumors of tobacco smokers. This signature displayed a weak association with proton and helium exposures, seen in 2 cell lines (Supp. Figure 4B). Certain features of ID83B and ID83D spectra are similar to the indel mutational signature provoked by ionizing photon radiation, reported in a recent preprint, where clonal organoids from mouse and human cells were irradiated^45^). Overall, because the association of ID83B does not appear specific to cell lines (i.e. there are no tissue-specific nor genetic background-specific effects), this suggests an universal indel mutational footprint of small deletions caused by particle-based radiation in human cells. However, due to additional indel signatures associated, we do not rule out that different cell types and/or genetic background might be differentially affected by different radiation types generating indel changes.. The activities of the indel signatures correlated with activities of some SNV signatures across our cohort (Supp. Figure 6), suggesting possible mechanistic links. In particular, SBS96D activity is significantly positively correlated with ID83C exposure, further supporting the idea that these signatures represent features of MMR deficiency.

### Mutational spectra of structural variation in irradiated cells

Large structural variants (SVs, also referred to as rearrangements) were grouped into four broad categories: large deletion (DEL), large insertions (INS), tandem duplications (DUP) and inversions (INV). No particular SV category has been found to be the most prevalent across the cell lines and across the cell lines/conditions (Figure 5A, Supp. Figure 1).

**Figure 5.**
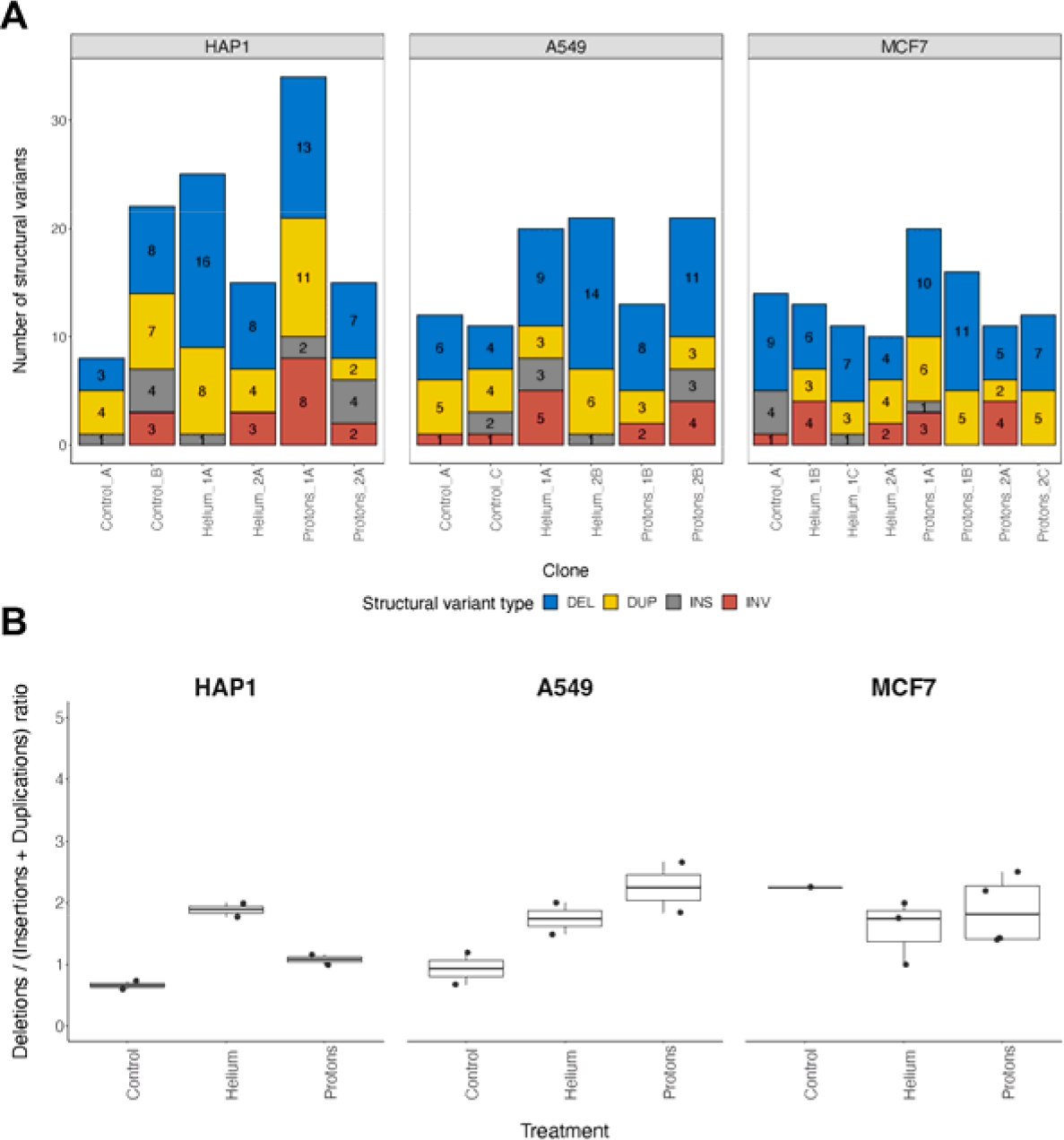
Structural variation in irradiated human cell lines. **A.** Distribution of the number of structural variants (SVs), decomposed into large deletions, large insertions, duplications and inversions, per clone in each of the three analyzed cell lines. **B.** Ratio of all large deletions divided by all large insertions and duplications, pooled by treatment for each cell line individually.

The rarest event type – insertions – is similarly observed across the cell lines and clones; their low frequency may however stem from technical difficulties in calling insertion variants from short-read WGS, resulting in many undetected variants. More generally, in our experiments, we observed globally a low number of SVs of any type, suggesting that the impact of the proton and alpha ray treatment on rearrangement rates in human cells may not be remarkable, at least in the treatment regimen applied here. Irradiated clones are not highly enriched in SVs when compared to control cells, although helium and proton-treated clones from all cell lines have higher numbers of large deletions compared to the untreated clones (Supp. Figure 1). We also note a substantial variability between individual clones in SV burden.

We also investigated another quantitative measure of the identified structural variation: the distributions of SV sizes, as commonly done in cancer genome analyses^27, 29^. Again, we asked if there is a difference between two types of radiation treatments, also comparing them with the non-irradiated controls. Because insertions were rarely detected, here we considered only deletions, duplications and inversions. While the distribution of SV lengths was largely not significantly different across the conditions, nevertheless, we observe some cell line specific trends. In HAP1, duplications are smallest in unirradiated clones and largest in proton treated clones, with a significant difference in length between the two (Supp. Figure 7, p=0.046). In A549 clones, on the other hand, the opposite is true: duplications in untreated clones are longest and significantly different from helium and proton clones (Supp. Figure 7, p=0.04 and p=0.026 respectively). Deletions and inversions are similar in size between all cell lines and treatment groups (Supp Figure 7). For inversions, the sample sizes are too small to ascertain any trends due to technical challenges of detection in short-read data. We observed a trend for deletions that is consistent across all the cell lines. In particular, proton radiation generated the smallest deletion events, followed by alpha, and finally longest deletions were observed in untreated controls (Supp. Figure 7). Inversions behaved similarly: we also observed that treated clones, whatever the radiation type, contain inversions smaller than those in non-treated samples. Moreover, there is also a trend toward longer inversions resulting from helium compared to protons (Supp. Figure 7). Overall, proton radiation generated SVs of shorter lengths than alpha rays or than the default mutagenesis in untreated cells, suggesting a different mode of DNA damage and/or repair thereof following proton exposure in various human cell types.

Finally, we investigated the enrichment of deletion SVs compared to insertion and duplication SVs (the latter two were considered to be similar and merged for this analysis). As previously reported for ionizing radiation-associated second malignancies^14^, the ratio of deletions to insertions increases in our particle-irradiated human cell lines when pooled together, and the trend stems from HAP1 and A549 clones (Figure 5B, Supp. Figure 1). Overall, different types of ionizing radiation, either X-rays or gamma as commonly employed for cancer therapy^13, 14^, or protons and helium/alpha particles as tested here, exhibit an overall similar spectrum of increase in deletion SVs relative to insertion/duplication SVs. When comparing this SV analysis to the indel analysis, considering the small deletion to insertion ratio, we observe a similar trend towards higher ratios in proton-treated clones compared to untreated (Supp. Figure 1 p=0.055 for small indels, p=0.082 for SVs). For large insertions and deletions though, we also observe a higher ratio in helium treated clones compared to control (Supp. Figure 1, unadjusted p=0.014), a trend which retains the direction but does not reach significance in the small indel ratio.

### Regional mutation rate with respect to replication time and gene activity

DNA replication timing (RT) is correlated with regional variability in somatic mutation rates in human tissues. There is an increased mutation rate in late-replicating heterochromatic regions of the genome, because DNA repair mechanisms preferentially protect the early-replicating euchromatic, gene-rich regions^47, 48^. We were interested if there is an enrichment of radiation-associated mutation rates in regions with a specific RT.

Overall SNV mutation rates were, as expected, associated with RT in all 3 cell lines (Supplementary Figure 8), with reduced rates in early replicating regions. The SVs observed did not display a significant association but weakly trended towards the converse association (Supplementary Figure 8); we note the modest number of SVs make it more difficult to detect associations. However, with respect to radiation treatment, we did not observe a a significant change of correlation between SNV or SV density and RT bins between treatments in any cell line (Supp Figure 8), suggesting that exposure to particle radiation does not impact the large-scale distribution of mutations along the RT domains in the genome.

Next, we considered differences in mutation rates at the gene scale by investigating the distribution of SNV and SV mutations across regions harboring various levels of gene expression (Supp Figure 8). Expectedly, higher gene expression was associated with lower SNV rates; we note a nonsignificant opposite trend for the SVs (Supp Figure 8). However there was no consistent effect of radiation treatment on the association between gene expression and SNV or SV burden across the three cell lines (we note some effect in SNV mutations that appears particular to the MCF7 cell line, and so tissue-specific effects or genetic background-specific effects cannot be ruled out). These results indicate that particle radiation treatment does not preferentially increase or decrease relative mutation rates in transcriptionally active regions of the genome.

### Clustered mutation processes due to irradiation

Next, we considered patterns of mutation clusters, which can be particularly informative about various mutational processes in human cells^28^, most prominently with regard to APOBEC enzyme DNA damaging activity^49–51^ and also error-prone DNA polymerase usage^28^. Operationally, we defined a mutation cluster as a set of point mutations at a pairwise intermutational distance lower than 1Kb; given the overall modest mutation burden in our genomes, this threshold is unlikely to result in artefactual (false-positive) mutation clusters. In the A549 cell line, 102 mutations were clustered in control clones, 171 in helium-irradiated clones and 143 in proton-irradiated samples. In the HAP1 cell lines, a total of 88 mutations were clustered in control clones, 178 in helium-irradiated clones and 255 in proton-irradiated samples. Finally, in the MCF7 cell line, we detected 130 mutations in the control clone, 325 clustered mutations in the helium-irradiated clones and 370 in the proton-irradiated clones (Supp. Figure 9A).

We detected more clustered mutations in radiation conditions compared to non-radiation conditions in all cell lines, indicating that alpha and proton radiation generates mutation clusters in human cells (Supp. Figure 9A). This pattern of mutagenesis is consistent with many reports of ionizing radiation damaging DNA in a clustered manner^52–54^. With respect to types of radiation, more clustered mutations are found in helium clones compared to proton clones in all cell lines, but the trend is particularly salient in HAP1 clones (Supp. Figure 9A).

We next asked if the clustered mutations, depending on the cell line and the radiation type, were found at closer distances. In HAP1 cells, clustered mutations were spaced significantly further apart for both helium and proton radiations compared to the control clones, with the helium group displaying the largest inter-mutational distance (Figure 6A). The trend is however not seen in the other cell lines, with all treatment groups having similarly spaced mutations, despite differences in mutation counts (Figure 6A).

**Figure 6.**
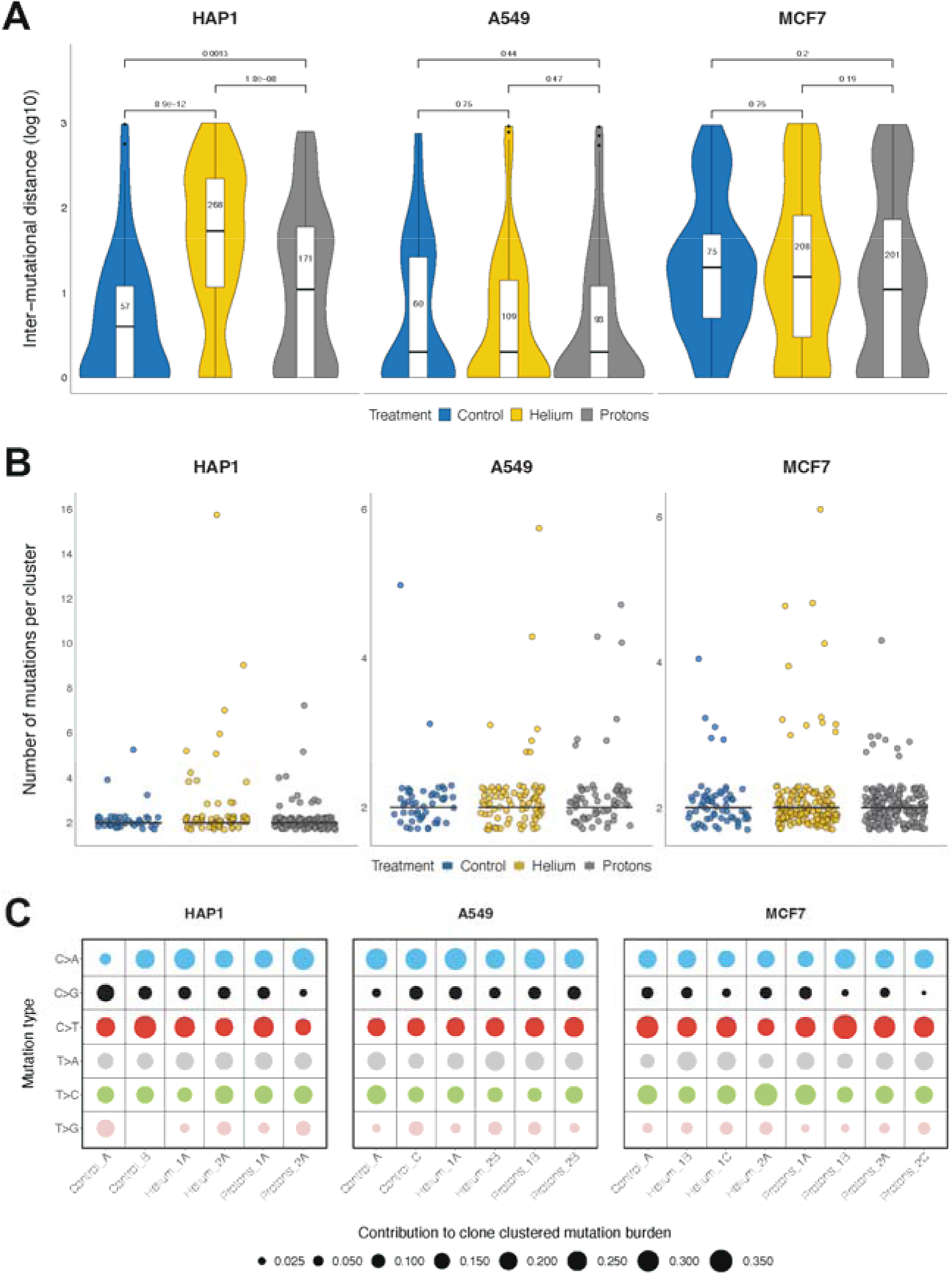
Clustered point mutations associated with irradiation. **A.** Distribution of the log10-transformed genomic distance between point mutations that were identified as being clustered together (see Methods for the definition of a mutational cluster); note the distance between two unique mutations is only counted once. The number assigned to each treatment denotes the total number of clustered mutations identified, whilst the black horizontal line represents the median inter-mutation distance. *P*-values shown were computed using the *ggpubr* R package v0.4.0 (t-test mean comparison of distances). **B.** Distribution of point mutation cluster sizes, pooled by treatment for each cell line individually. C. Break-down of clustered point mutations into the 6 mutation classes, relative contributions per clone in each cell line. Empty tiles signify the absence of mutation clusters in the given clone.

We were interested in the arrangement of the clustered mutations, *i.e.* do they form a lot of small clusters or do they form a few large (multi-mutation) clusters, the latter case corresponding to a kataegis-like phenomenon^49^ and the former to an omikli-like phenomenon^51^. We applied a graph-theory approach to quantify this (see Methods). Considering large clusters as connected components of size higher than 5, we identified one large cluster in A549/helium, four in HAP1/helium, one in MCF7/helium and one in HAP1/proton. However there were no large clusters in the untreated cells, suggesting mutation showers can be generated by ionizing radiation (Figure 6B). Of note, the majority of clustered mutations, both in the irradiated and in the untreated conditions, were in smaller, *omikli*-like clusters; for most cell lines and conditions, over 90% of clusters had a size of 2 (Figure 6B).

Finally, we were interested in the mutational spectrum of the clustered SNV mutations, which can identify or rule out a mechanism underlying the clusters. Here, because of the relatively modest number of SNVs, we considered the 6 relative contribution mutation type-spectrum and compared it with the spectrum for all SNVs (Figure 3A).

The clustered SNV spectrum (Figure 6C, Supp. Figure 9B) differed from the general SNV spectrum in that the clustered spectrum had a substantially lower proportion of C>A changes (largely due to oxidative stress during cell culture), and a somewhat lower proportion of T>C transitions (which can result from multiple causes). The proportion of C>G and T>G changes is similar between the clustered and unclustered. C>T and T>A changes are slightly more prevalent in clustered mutations compared to the total, although variation between clones exist. The lack of high C>G enrichment in clustered mutations, as well as the lack of TCN > G/T mutations in the 96-class spectrum (Supp. Figure 12, Supp. Table 2) suggests that the activity of the APOBEC3 cytidine deaminase is likely not responsible for the clustered mutation burden in our experiments.

Overall, this suggests that the clustering of mutations often results from exposure to particle radiation, that the underlying mechanism is unlikely to be APOBEC-related, that a process with a similar mutation spectrum is active (albeit at much lower levels) also in unirradiated cells,, and that it may occasionally generate multi event, kataegis-like mutation clusters.

## Discussion

In this study, we analyzed genomes of three human cancer cell lines irradiated with two different types of GCR – proton and helium fluxes – in order to characterize the spectra of DNA alterations that GCR would produce in different human tissues. Overall burden of SNV mutations, indels and SV events was not markedly different between treated and untreated cells. This suggests that GCR exposure – at least in the regime applied in our experiments – is not grossly mutagenic (see below for further considerations).

However, there were certain differences in mutation spectra and distributions between GCR-treated and untreated cells. For instance, in agreement with a recent study analysing ionizing radiation-associated tumors^14^, in indels we observed an enrichment of deletions compared to insertions, and this was even more the case for large (SV) events (Figure 4C, Figure 5B). This phenomenon was also observed in a recent study that reported a 3.6-fold increase of small deletion burden when comparing radiotherapy treated glioma patients to untreated glioma patients^13^. We note that tumor radiotherapy usually employs X rays or gamma rays, rather than the protons and alpha particles tested here, and so this enrichment of deletion mutations appears broadly independent of the type of IR applied to cells.

We found that GCR generates point mutations that are more often clustered than those found in non-irradiated samples. This is in line with recent work suggesting clustered mutations to be more specific indicators of certain types of mutagenic processes^28, 55, 56^ than the general, genome-wide mutation signatures. The SNV spectra of the clustered mutational distributions generated by helium and proton radiation exposure do not indicate a major role of the known agents commonly generating clustered mutations in human cells: the APOBEC3A enzyme and the translesion synthesis DNA polymerase eta (POLH)^28, 55^. Mechanisms would be clarified with additional experiments in future work.

We were also interested in the SNV trinucleotide mutational signatures that helium and proton GCR can generate, and whether known biological mechanisms can be linked to those footprints. While most observed SNV signatures were consistent across all treated and untreated samples, e.g. reactive oxygen species mutagenesis was noted universally, some samples exhibited signatures consistent with MMR deficiency (this was however unlikely to have resulted from, or have been selected by irradiation, because we detected it also in a control sample). Interestingly, we observed a modest enrichment of certain SNV signatures in both irradiation conditions : a C>A signature linked with oxidative stress, and an T>A rich SBS signature, resembling previous signatures of aristolochic acid and some chemotherapy treatments^49, 50^, suggesting, speculatively, that particle radiation may generate DNA damages resembling bulky adducts in some cell types.

Similar to SNVs, some indel signatures were radiation-associated, being consistently enriched (with modest effects) exposed samples, thus suggesting a likely general effect over tissues for indel-generating mechanisms. The indel signatures bore certain similarities with indel signatures previously reported for photon ionizing radiation^30, 45^; additional data appears required to gain statistical power to establish to what extent the indel signatures indeed differ between photons and particle rays, providing more insight into mechanisms.

Regading non-ionizing radiation, the solar radiation-generated SNV signatures identified previously^15, 31^ (presumably generated by the UV radiation) were observed at low intensities in our irradiated samples, likely stemming from imprecisions during statistical analysis. This is consistent with prior knowledge that ionizing cosmic radiation affects the DNA differently than the non-ionizing UV radiation^56^.

Our study also has several limitations. For instance, some components of GCR were not considered in this study, in particular the heavy atom nuclei (often referred to as “HZE ions”), highly energetic particles likely originating from supernova explosions. Despite their small proportion in cosmic rays, their biological impact might be large, and their mutagenic effects on genome stability remain to be studied.

Next, we considered only a single dose of radiation that resulted in a ∼40% to 50% reduction of cell viability assessed by the colony formation assay, and a fractionation regime of four total fractions (with each fraction given every four days) exposures of this dose. The dose studied here had rather strong biological effects on cell viability, and does not correspond to what would be encountered in space travel (where any vessel would need to be shielded to prevent such exposures that would have very deleterious acute effects on the organism). These exposures were chosen as an experimental setup, with the rationale that dosages with notable effects on cell viability would probably be well sufficient to observe the mutagenic effects of GCR, if any. Since a gross increase in mutation rates was not observed, we infer that GCR exposures are not highly mutagenic. However, we cannot rule out the (less parsimonious) scenario where lower doses of GCR than those employed here might be more mutagenic e.g. by failing to trigger cell cycle checkpoints, thus ‘slipping under the radar’ of the mechanisms protecting genome integrity and introducing mutations.

Moreover, given that for various mutation patterns we observed responses that appear specific to one cell line, this suggests there may be tissue-specific and/or genetic background specific responses (these scenarios cannot be distinguished from our data). Additional experiments would be needed on different cell lines or other experimental models to ascertain tissue-specific responses to various radiation types. In a related vein, this study was performed on cancer cell lines, and mutational responses might be different in healthy, noncancerous cells, which remains to be investigated in future work.

In conclusion, our data overall suggests that particle ray components of GCR are not overtly mutagenic to human cells, with very modest effects on point mutation spectra, and with some effects on SV distributions, and on indels and particularly on clustered mutation spectra. However, the intermittent exposure regime that we employed is different from longer-term, chronic exposures to GCR expected e.g. during space flight. Even modest increases in mutation rates under chronic GCR exposures might have detrimental effects on cancer risk, and possibly neurodegeneration and reproductive health and on genetic disease incidence in the progeny. Therefore we highlight the necessity of further experimental work on cell and animal models, using longer-term exposures to various components of galactic cosmic rays to ascertain their effects on the stability of the human genome.

## Methods

### Cell lines

We used three human cell lines: A549, HAP1 and MCF7. The lung adenocarcinoma cell line A549 (CCL-185) and the breast adenocarcinoma cell line MCF7 (HTB-22) were purchased from ATCC (the American Type Culture Collection, Manassas, VA). The near-haploid cell line derived from the KBM-7 cell line HAP1 (C859) was purchased from Horizon (Carle Place, NY). All cell lines were cultured and maintained according to recommended protocols. All the cell lines were authenticated by STR profiling by the corresponding repositories.

All cells were grown at 37°C in a 5% CO_2_ humidified incubator; the A549 cells were grown in Eagle’s Minimum Essential Medium (MEM), MCF7 in Dulbecco modified MEM, and HAP-1 cells in Iscove’s modified Dulbecco’s medium (IMDM). All the media were supplemented with 10% heat-inactivated (56°C, 30 min) fetal bovine serum (FBS), and 100 U/ml penicillin and 100 mg/ml streptomycin (all from Sigma–Aldrich Corp).

### Cell irradiation, culture and clones isolation

A clonogenic assay^57^ pilot experiment was conducted to assess the dose that resulted in 40-50% lethality (i.e., 50-60% survival) to 11 MeV helium ions (LET = 65 keV/µm) or 5.4 MeV protons (LET = 10 keV/µm), which was 0.5 and 1 Gy, respectively (Figure 1A,B). Cells for the experiment were then exposed to four total fractions delivered every four days. Cells were collected four days after the last fraction. For each sample, half of the cells were frozen at -80 C while the other half was re-plated until confluency. The cells were then collected, separated by vigorously passing through a 21 G syringe and 3 ul/ml of DAPI was added to the solution for cell sorting at the BD Influx cell sorter. For each cell line, single cells were sorted in two 96-well plates and incubated in 250 ul of medium supplemented with 20% FBS. Once colonies were visible, we picked ∼30 colonies and re-plated each one in an individual well of a 12-well plate in the case of A549 and MCF7 and in 48-well plate in the case of HAP1 cells up to confluence. The cells were then collected and half frozen at -80C while the other half was replated in 6-well plates and then in 100-mm Petri dish until confluence. DNA was extracted with PureLink Genomic DNA kit (Invitrogen), frozen and shipped to the sequencing centre.

### Cell irradiations at the track segment irradiation platform

The 5.5 MV Singletron accelerator at the Radiological Research Accelerator Facility (RARAF) served as a source of energetic protons and alpha particles. The accelerator was operated at the maximum terminal voltage generating beams with nominal energies of 5.5 MeV and 11 MeV for protons and alphas, respectively. Cells were irradiated at the so called “track segment” irradiation platform whose name indicates the traversal of thin samples (typically cell monolayers) by a short segment of the ion’s trajectory resulting in a small variation of the linear energy transfer (LET) throughout the sample. It is valid therefore to assume that radiation doses at the track segment platform are delivered by mono-LET beams. The LET values of the applied beams were 10 keV/µm for protons and 65 keV/µm for alphas. Detailed description of the track segment irradiation platform and its operational principles can be found elsewhere^1–4^. We will state here only the main features of the irradiation protocol. The protocol can be generally divided in two parts. First, beam characterization and dosimetry were performed with the use of different detectors mounted on the metal wheel that rotates over the beam exit aperture. The aims of this step were to verify the energy and the LET of the beam, to check the uniformity of the irradiation field covering the 6 mm x 20 mm beam exit aperture (2.9 µm thick havar foil) and, finally, to calibrate the online beam monitor in terms of the absorbed dose delivered to the sample. In the second step, the dosimetry wheel was replaced with the sample-carrying wheel that can accommodate up to 20 custom made dishes. The dishes were manufactured by gluing 6 µm thick mylar foil over metal rings having a diameter of 5 cm. The mylar foil serves as the bottom of the dish on which the cells are attached, allowing thus the ions to penetrate through and reach the cells without losing much of their energy. The in-house developed software drives the stepper motor and controls the movement of sample dishes over the beam. A desired dose was delivered to the sample by exposing it to the beam until the appropriate number of monitor counts was reached according to dosimetry measurements performed during the first part of the irradiation protocol.

### Irradiations of the cells for mutational analysis and clone isolations

The obtained survival curves were fitted with the linear quadratic (LQ) model. Doses of 1 Gy and 0.5 Gy were selected for investigating the mutational signatures of protons and alpha particles, respectively. According to the LQ fits, the selected doses resulted in preservation of between 50% and 60% clonogenic capacity for all cell lines. Four fractions of the same dose were delivered to the cells every 96 hours to amplify the number of mutations in surviving cells. Between the fractions, the cells were not removed from the mylar dishes. To prevent the cells from overgrowing the dish size before the end of the irradiations, the initial number of plated cells was < 1000.

### Whole-genome sequencing

Total genomic DNA was extracted from pelleted cells using PureLink™ Genomic DNA Mini Kit (K182001, Thermo Fisher Scientific). DNA was sequenced on NovaSeq 6000, 150 nt paired-end mode. Reads were aligned to the reference human genome Hg38 (GRCh38.d1.vd1) using BWA v0.7.17^58^. GATK Base Quality Score Recalibration was applied to adjust quality scores assigned during the sequencing process^59^. The average coverage of the various samples was 18.5X to 44.8X.

### Variant detection

Strelka2 variant calling algorithm v2.9.10 was used to detect point mutations and small insertions and deletions^60^. Strelka2 was launched in the joint genotyping mode: briefly, mutations were called for each clone individually (in one batch), but genotyped jointly across all clones belonging to a given cell line. . Mutations were then annotated using annovar, function table_annovar.pl^61^ with the gnomAD genome database v2.1.1 allele frequencies^62^ This allowed us to identify mutations shared by all clones (which were acquired prior to treatment) as well as discrete mutations acquired by only one clone during treatment. To arrive at the final set of mutations used in our analyses, a number of filters were applied: 1) variants passed both sample and call filters from Strelka2, 2) variants were found in uniquely mapping region of the genome (based on the Umap k50 mappability tracks), 3) variants were either not present in the gnomAD database or found at frequencies lower than 0.1%, 4) variants were genotyped in only one of the cell line clones, 5) variants were not multiallelic and 6) there were 2 or more variant supporting reads in the genotyped clone. For small insertions and deletions, we also imposed a genotype quality equal or higher than 10 and removed variants where more than two other clones have one or more variant supporting reads and variants where more than one other clone has two or more supporting reads, in order to remove putatively germline mutations only genotyped in one clone by Strelka2.

To call structural variants, i.e. large insertions and deletions, duplications and inversions, we used the Manta v1.6.0 tool in the joint calling mode^63^, in the same manner we used Strelka2. For inversions, Manta returns a list of breakpoints that describe the inversions it detected. To transform those breakpoints into inversions, we applied the convertInversion.py python script provided by the Manta developers (https://github.com/Illumina/manta). Filters 1-5 described above were also used to filter the structural variant calls; further, we required variants to have 2 or more supporting split-reads in the genotyped clone, with no requirement for the number of supporting spanning reads.

### Mutation clustering

Point mutations were considered to be clustered together if they appear at positions with a genomic distance lower than 1kb; such mutations would be classified asomikli (“mutation fog”)^51^. To consider large, multi-mutation clusters resembling a kataegis (mutation shower) event^49, 64^, we used an approach based on graph theory. Roughly, we consider each mutation to be a node of the graph, and edges connect two nodes if the distance between corresponding mutations is lower than 1kb. Then, we computed the components of the graph, that corresponds to a connected sub-graph, and subsequently the size of the components, *i.e.* how many nodes or mutations are present in the component.

### Extraction of mutational signatures

We pooled our samples (21 clones) with two already published datasets in order to help NMF converge and in order to compare our datasets to signatures of DNA damage and of various mutagens. We first pooled our data with the mutations reported in the Zou *et al.* study^39^, which analysed 38 clones generated by CRISPR-Cas9 knockouts of 9 DNA repair/replicative pathway genes (ΔOGG1, ΔUNG, ΔEXO1, ΔRNF168, ΔMLH1, ΔMSH2, ΔMSH6, ΔPMS1, and ΔPMS2) which generated mutational signatures in human induced pluripotent stem cells (note that there were additional gene knockouts which did not produce mutational signatures). We then pooled the data with the mutations reported in the Kucab *et al.* study^31^, which reported 153 clones (treated with 53 environmental agents) yielding a mutational signature (note there were additional samples in the study without a prominent mutation pattern, which were not included here), as well as 2 clones treated with gamma radiation (with no identified signature).This pooled dataset of 214 individual clones was used for SNV and indel signature extraction. . In order to extract SNV signatures generated by our radiation treatments, we applied SigProfiler ExtractorR v1.1.16 algorithm^15^ with default settings on the set of detected mutations, classified into the 96 SNV categories. The suggested solution contained 10 extracted signatures with high stability and low reconstruction errorExtracted signature spectra were compared and assigned to a known PCAWG “SBS” signature set^15^ (COSMIC_v3.3.1_SBS_GRCh38 from https://cancer.sanger.ac.uk/signatures/downloads/) if the cosine similarity between the two mutational profiles was at least 0.85. Further, all signatures were decomposed into the same set of PCAWG signatures using the python SigProfiler Assignment v0.0.24 tool (https://github.com/AlexandrovLab/SigProfilerAssignment), function decompose_fit.

In order to determine the extent to which the addition of Zou et al. and Kucab et al. samples influences the extracted SNV signature spectra and their respective exposures in our dataset, we implemented a subsampling approach whereby half of each external dataset is randomly removed thrice from the original dataset and the NMF algorithm is run on the remaining 117 samples (19 from Zou et al., 77 from Kucab et al. and 21 from our dataset), once for every subsampled set. Then, mutational signature spectra and exposures in our samples are compared to the original set.

For indel signatures, we used the SigProfiler Matrix Generator for R v1.2.4 tool to correctly classify the observed indels into one of the 83 categories defined in Alexandrov *et al.*^15^. Then, the SigProfiler ExtractorR v1.1.16 tool was launched on the same set of 214 samples from Kucab *et al.*, Zou *et al.*, and our experiments, using the generated mutation matrices. We considered the optimal solution given by the tool, resulting in a total set of 4 extracted indel signatures with high stability and low reconstruction error. As for the SNV signatures, we compared and decomposed our extracted spectra into the known PCAWG “ID” signature set^15^ (COSMIC_v3.3_ID_GRCh37.txt from https://cancer.sanger.ac.uk/signatures/downloads/).

### Regional enrichment analysis

We performed a regional enrichment analysis in order to estimate if the mutations detected in the irradiated clones were located in regions with a specific epigenetic pattern compared to the mutations detected in the non-treated clones. We tested three genomic features: the replication timing, the DNA repair histone mark H3K36me3 and the gene expression levels based on RNA-seq data. Replication timing bins, genomic bins extracted from chromatin mark H3K36me3, and regions based on variable gene expression were computed as in Supek and Lehner^28^. To detect significant association of mutations in specific genomic regions, we fitted a negative binomial regression using the glm.nb function from the MASS R package, on the counts of mutations per bins, controlling for the 3-nucleotide context and type of mutation in the case of SNV, as in Supek and Lehner^28^. The regional enrichment analysis source code is implemented using Nextflow^65^ and is available on the GitHub platform (https://github.com/tdelhomme/RegionalEnrichment-nf/).

### Statistical analyses

All the statistical analyses were performed using R software version 3.6.0. VariantAnnotation^66^ R package v1.32 was used to read the VCF files. The R packages BSgenome.Hsapiens.UCSC.hg38 v1.4.1 and BSgenome.Hsapiens.UCSC.hg19 v1.4.0 were used to load the human genome references inside R. MutationalPatterns^67^ R package v1.12 was used to generate the trinucleotide contexts. Ggraph v2.0.5 and Igraph v1.2.6 R packages were used to identify the cluster components. The statistical tests used to derive the p-values within some figures are described in the figure legends.

In order to systematically compare the mutation burdens and mutation signature exposures between all cell lines and treatments, we implemented a randomization test. Taking every mutation type (in the case of mutation burden comparison) and every signature (in the case of exposure comparisons), we randomly shuffled the clone labels (n=21) 100.000 times. Then, we calculated the real mean mutation burden or exposure for each cell line (stratified by treatment and pooled) and for each treatment (stratified by cell line and pooled) and the mean from each shuffling iteration, resulting in 1 real mean and 100.000 randomised means for each comparison. Then, we calculated the pairwise difference between the mean of all cell lines and treatments (stratified or pooled) and compared the real mean difference to the distribution of randomised mean differences. For each such comparison, we calculated two empirical p-values^68^, where p=(r+1)/(n+1), r is the number of randomised differences >= or <= than the real mean difference and n is the total number of iterations (100.000). Then, all p-values from all comparisons across SNV signatures, ID signatures and mutation burdens were corrected (separately for each category) to account for the multiple comparisons problem using the p.adjust function from R core package stats v3.6.0, method “BH”/”fdr”^69^. Comparisons with an original p-value of 0.1 or lower were reported with their corrected values.

## Data Availability

All Variant calling files (VCFs) derived from the WGS data generated in this study (see Methods for details) are available on FigShare repository using the following link: https://figshare.com/s/4f071118f2fde3f00a26

Relevant source code for statistical analyses from this study is provided on GitHub at the following link: https://github.com/maia-munteanu/Radiation_paper_2023

## Acknowledgements

Work in the F.S. lab was supported by the ERC Starting Grant “HYPER-INSIGHT” (grant number 757700 to F.S.); the Horizon2020 RIA project “DECIDER” (grant number 965193 to F.S.); the Spanish Ministry of Science, Education and Universities project “REPAIRSCAPE” (grant number PID2020-118795GB-I00 to F.S.); the State Agency for Research of the Ministry of Science and Innovation - Severo Ochoa Centre of Excellence Award (grant number CEX2019-000913-S to IRB Barcelona); and the CERCA Generalitat de Catalunya funds (to IRB Barcelona). We thank Jurica Levatić for a literature review, input on irradiation protocols and on stimulating discussions motivating the lab’s work on mutagenesis due to space radiation. T.D. was supported by a Juan de la Cierva fellowship from the Spanish Ministry of Science, Education and Universities. M.B. acknowledges support from the National Cancer Institute (NCI) grant U01CA236554.

## Author Contributions

T.D. and M.M. performed all computational analyses. M.B., V.G. and J.B. performed experiments. F.S. supervised the study. All authors drafted and edited the manuscript.

## References

1. Larose, T. L. Tumors in Space: Preparation for Spaceflight. Preparation of Space Experiments (IntechOpen, 2020). doi:10.5772/intechopen.93465.

2. Jastrow, R. Definition of Air Space. in First Colloquium on the Law of Outer Space (eds. Haley, A. G. & Heinrich, W.) 82–82 (Springer, 1959). doi:10.1007/978-3-7091-4414-5_16.

3. 100km Altitude Boundary for Astronautics | World Air Sports Federation. https://www.fai.org/page/icare-boundary (2017).

4. McDowell, J. C. The edge of space: Revisiting the Karman Line. Acta Astronaut. 151, 668–677 (2018).

5. Thirsk, R., Kuipers, A., Mukai, C. & Williams, D. The space-flight environment: the International Space Station and beyond. CMAJ Can. Med. Assoc. J. 180, 1216–1220 (2009).

6. Panesar, S. S. & Ashkan, K. Surgery in space. Br. J. Surg. 105, 1234–1243 (2018).

7. Hellweg, C. E. & Baumstark-Khan, C. Getting ready for the manned mission to Mars: the astronauts’ risk from space radiation. Naturwissenschaften 94, 517–526 (2007).

8. Cucinotta, F. A. Space Radiation Risks for Astronauts on Multiple International Space Station Missions. PLOS ONE 9, e96099 (2014).

9. Nikiforov, Y. E. Thyroid carcinoma: molecular pathways and therapeutic targets. Mod. Pathol. Off. J. U. S. Can. Acad. Pathol. Inc 21 **Suppl 2**, S37–43 (2008).

10. Morton, L. M. et al. Radiation-related genomic profile of papillary thyroid carcinoma after the Chernobyl accident. Science 372, eabg2538 (2021).

11. Brenner, A. V. et al. I-131 dose response for incident thyroid cancers in Ukraine related to the Chornobyl accident. Environ. Health Perspect. 119, 933–939 (2011).

12. Ozasa, K. Epidemiological research on radiation-induced cancer in atomic bomb survivors. J. Radiat. Res. (Tokyo) 57 **Suppl 1**, i112–i117 (2016).

13. Kocakavuk, E. et al. Radiotherapy is associated with a deletion signature that contributes to poor outcomes in patients with cancer. Nat. Genet. 53, 1088–1096 (2021).

14. Behjati, S. et al. Mutational signatures of ionizing radiation in second malignancies. Nat. Commun. 7, 12605 (2016).

15. Alexandrov, L. B. et al. The repertoire of mutational signatures in human cancer. Nature 578, 94–101 (2020).

16. Berrington de González, A. & Darby, S. Risk of cancer from diagnostic X-rays: estimates for the UK and 14 other countries. Lancet Lond. Engl. 363, 345–351 (2004).

17. IARC. Ionizing Radiation, Part 1: X- and Gamma (γ)-Radiation, and Neutrons.

18. Hughes, J. R. & Parsons, J. L. FLASH Radiotherapy: Current Knowledge and Future Insights Using Proton-Beam Therapy. Int. J. Mol. Sci. 21, 6492 (2020).

19. Yuan, T.-Z., Zhan, Z.-J. & Qian, C.-N. New frontiers in proton therapy: applications in cancers. Cancer Commun. Lond. Engl. 39, 61 (2019).

20. Li, M. et al. Clinical Efficacy and Safety of Proton and Carbon Ion Radiotherapy for Prostate Cancer: A Systematic Review and Meta-Analysis. Front. Oncol. 11, 709530 (2021).

21. Hu, M., Jiang, L., Cui, X., Zhang, J. & Yu, J. Proton beam therapy for cancer in the era of precision medicine. J. Hematol. Oncol.J Hematol Oncol 11, 136 (2018).

22. Vogel, J. et al. Proton therapy for pediatric head and neck malignancies. Pediatr. Blood Cancer 65, (2018).

23. Jain, V. et al. Predicted Secondary Malignancies following Proton versus Photon Radiation for Oropharyngeal Cancers. Int. J. Part. Ther. 6, 1–10 (2020).

24. Huang, J. & Mehta, M. Can proton therapy reduce radiation-related lymphopenia in glioblastoma? Neuro-Oncol. 23, 179–181 (2020).

25. Mohan, R. et al. Proton therapy reduces the likelihood of high-grade radiation-induced lymphopenia in glioblastoma patients: phase II randomized study of protons vs photons. Neuro-Oncol. 23, 284–294 (2021).

26. Alexandrov, L. B., Nik-Zainal, S., Wedge, D. C., Campbell, P. J. & Stratton, M. R. Deciphering signatures of mutational processes operative in human cancer. Cell Rep. 3, 246–259 (2013).

27. Li, Y. et al. Patterns of somatic structural variation in human cancer genomes. Nature 578, 112–121 (2020).

28. Supek, F. & Lehner, B. Clustered Mutation Signatures Reveal that Error-Prone DNA Repair Targets Mutations to Active Genes. Cell 170, 534–547.e23 (2017).

29. Degasperi, A. et al. Substitution mutational signatures in whole-genome-sequenced cancers in the UK population. Science 376, science.abl9283 (2022).

30. Pleasance, E. et al. Pan-cancer analysis of advanced patient tumors reveals interactions between therapy and genomic landscapes. *Nat*. Cancer 1, 452–468 (2020).

31. Kucab, J. E. et al. A Compendium of Mutational Signatures of Environmental Agents. Cell 177, 821–836.e16 (2019).

32. Saini, N. et al. The Impact of Environmental and Endogenous Damage on Somatic Mutation Load in Human Skin Fibroblasts. PLOS Genet. 12, e1006385 (2016).

33. Franco, I. et al. Whole genome DNA sequencing provides an atlas of somatic mutagenesis in healthy human cells and identifies a tumor-prone cell type. Genome Biol. 20, 285 (2019).

34. Davidson, P. R., Sherborne, A. L., Taylor, B., Nakamura, A. O. & Nakamura, J. L. A pooled mutational analysis identifies ionizing radiation-associated mutational signatures conserved between mouse and human malignancies. Sci. Rep. 7, 7645 (2017).

35. Sherborne, A. L. et al. Mutational analysis of ionizing radiation-induced neoplasms. Cell Rep. 12, 1915–1926 (2015).

36. Volkova, N. V. et al. Mutational signatures are jointly shaped by DNA damage and repair. Nat. Commun. 11, 2169 (2020).

37. MCF7 - HTB-22 | ATCC. https://www.atcc.org/products/htb-22.

38. Olbrich, T. et al. A p53-dependent response limits the viability of mammalian haploid cells. Proc. Natl. Acad. Sci. U. S. A. 114, 9367–9372 (2017).

39. Zou, X. et al. A systematic CRISPR screen defines mutational mechanisms underpinning signatures caused by replication errors and endogenous DNA damage. *Nat*. Cancer 2, 643–657 (2021).

40. Costello, M. et al. Discovery and characterization of artifactual mutations in deep coverage targeted capture sequencing data due to oxidative DNA damage during sample preparation. Nucleic Acids Res. 41, e67 (2013).

41. Sekiguchi, M. & Tsuzuki, T. Oxidative nucleotide damage: consequences and prevention. Oncogene 21, 8895–8904 (2002).

42. Kawanishi, S., Hiraku, Y. & Oikawa, S. Mechanism of guanine-specific DNA damage by oxidative stress and its role in carcinogenesis and aging. Mutat. Res. 488, 65–76 (2001).

43. Wang, R. et al. OGG1-initiated base excision repair exacerbates oxidative stress-induced parthanatos. Cell Death Dis. 9, 1–15 (2018).

44. Mas-Ponte, D., McCullough, M. & Supek, F. Spectrum of DNA mismatch repair failures viewed through the lens of cancer genomics and implications for therapy. Clin. Sci. Lond. Engl. 1979 136, 383–404 (2022).

45. Youk, J. et al. Mutational impact and signature of ionizing radiation. 2021.01.12.426324 Preprint at https://doi.org/10.1101/2021.01.12.426324 (2021).

46. Nguyen, L., W. M. Martens, J., Van Hoeck, A. & Cuppen, E. Pan-cancer landscape of homologous recombination deficiency. Nat. Commun. 11, 5584 (2020).

47. Supek, F. & Lehner, B. Differential DNA mismatch repair underlies mutation rate variation across the human genome. Nature 521, 81–84 (2015).

48. Zheng, C. L. et al. Transcription restores DNA repair to heterochromatin, determining regional mutation rates in cancer genomes. Cell Rep. 9, 1228–1234 (2014).

49. Nik-Zainal, S. et al. The life history of 21 breast cancers. Cell 149, 994–1007 (2012).

50. Alexandrov, L. B. et al. Signatures of mutational processes in human cancer. Nature 500, 415–421 (2013).

51. Mas-Ponte, D. & Supek, F. DNA mismatch repair promotes APOBEC3-mediated diffuse hypermutation in human cancers. Nat. Genet. 52, 958–968 (2020).

52. Sage, E. & Shikazono, N. Radiation-induced clustered DNA lesions: Repair and mutagenesis. Free Radic. Biol. Med. 107, 125–135 (2017).

53. Sutherland, B. M., Bennett, P. V., Weinert, E., Sidorkina, O. & Laval, J. Frequencies and relative levels of clustered damages in DNA exposed to gamma rays in radioquenching vs. nonradioquenching conditions. Environ. Mol. Mutagen. 38, 159–165 (2001).

54. Hada, M. & Georgakilas, A. G. Formation of clustered DNA damage after high-LET irradiation: a review. J. Radiat. Res. (Tokyo) 49, 203–210 (2008).

55. Roberts, S. A. et al. An APOBEC cytidine deaminase mutagenesis pattern is widespread in human cancers. Nat. Genet. 45, 970–976 (2013).

56. Ravanat, J.-L. & Douki, T. UV and ionizing radiations induced DNA damage, differences and similarities. Radiat. Phys. Chem. 128, 92–102 (2016).

57. Puck, T. T. & Marcus, P. I. Action of x-rays on mammalian cells. J. Exp. Med. 103, 653–666 (1956).

58. Li, H. & Durbin, R. Fast and accurate short read alignment with Burrows-Wheeler transform. Bioinforma. Oxf. Engl. 25, 1754–1760 (2009).

59. McKenna, A. et al. The Genome Analysis Toolkit: a MapReduce framework for analyzing next-generation DNA sequencing data. Genome Res. 20, 1297–1303 (2010).

60. Kim, S. et al. Strelka2: fast and accurate calling of germline and somatic variants. Nat. Methods 15, 591–594 (2018).

61. Wang, K., Li, M. & Hakonarson, H. ANNOVAR: functional annotation of genetic variants from high-throughput sequencing data. Nucleic Acids Res. 38, e164 (2010).

62. Karczewski, K. J. et al. The mutational constraint spectrum quantified from variation in 141,456 humans. Nature 581, 434–443 (2020).

63. Chen, X. et al. Manta: rapid detection of structural variants and indels for germline and cancer sequencing applications. Bioinforma. Oxf. Engl. 32, 1220–1222 (2016).

64. Roberts, S. A. et al. Clustered mutations in yeast and in human cancers can arise from damaged long single-strand DNA regions. Mol. Cell 46, 424–435 (2012).

65. Di Tommaso, P. et al. Nextflow enables reproducible computational workflows. Nat. Biotechnol. 35, 316–319 (2017).

66. Obenchain, V. et al. VariantAnnotation: a Bioconductor package for exploration and annotation of genetic variants. Bioinforma. Oxf. Engl. 30, 2076–2078 (2014).

67. Blokzijl, F., Janssen, R., van Boxtel, R. & Cuppen, E. MutationalPatterns: comprehensive genome-wide analysis of mutational processes. Genome Med. 10, 33 (2018).

68. Davison, A. C. & Hinkley, D. V. Bootstrap Methods and their Application. (Cambridge University Press, 1997). doi:10.1017/CBO9780511802843.

69. Benjamini, Y. & Hochberg, Y. Controlling the False Discovery Rate: A Practical and Powerful Approach to Multiple Testing. J. R. Stat. Soc. Ser. B Methodol. 57, 289–300 (1995).

