## Supplementary Figures 1-9 for "Proton and alpha radiation-induced mutational profiles in human cells"

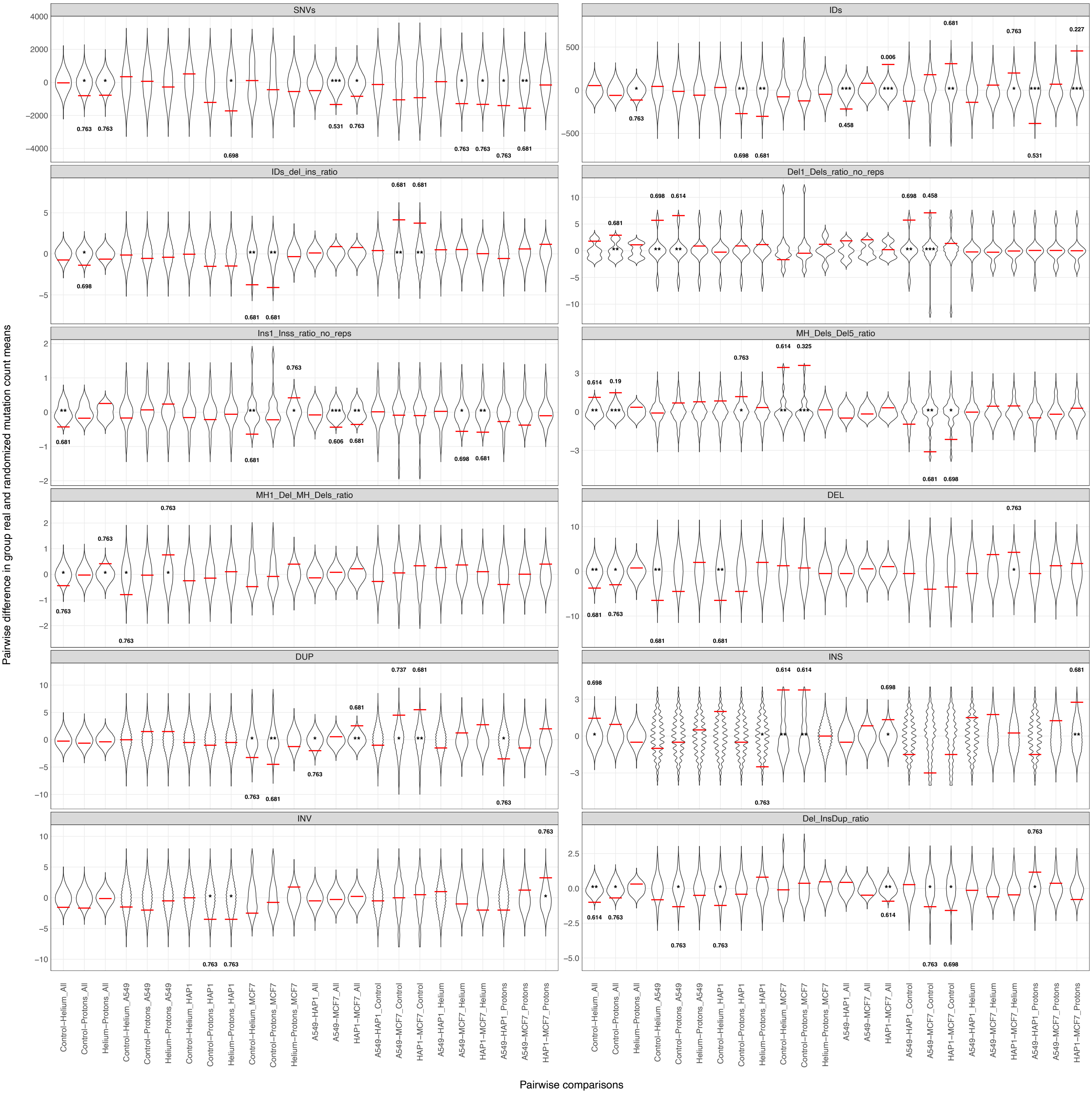


**Supp. Figure 1. Mutation burden randomization tests.** Pairwise differences in treatment/cell line group means (pooled or stratified by treatment or cell line) of mutation burdens (SNVs - point mutations; IDs - small insertions and deletions; IDs_del_ins_ratio - ratio of small deletions divided by small insertions; Del1_Dels_ratio_no_reps - ratio of 1 bp deletions divided by >1 bp small deletions excluding all variants at homopolymer repeats; Ins1_Inss_ratio_no_reps - ratio of 1bp insertions divided by >1 bp small insertions excluding all variants at homopolymer repeats; MH_Dels_Del5_ratio - ratio of deletions <5 bp with microhomology divided by deletions >=5 bp with microhomology; MH1_Del_MH_Dels_ratio - ratio of deletions with 1 bp microhomology divided by deletions with >=2 bp microhomology; DEL - large deletions; DUP - duplications; INS - large insertions; INV - inversions; Del_InsDup_ratio - ratio of large deletions divided by large insertions and duplications). Red horizontal lines represent the difference between real group means, whereas the violin distributions represent the differences between randomized group means. Unadjusted empirical p-values are marked using the following scale: (*) p <= 0.1, (**) p <= 0.05, (***) p <= 0.01. Q-values are printed for all comparisons with a p-value <= 0.1 on the side of the distribution where the real difference in mean is located (top - the real mean difference is higher than most points in the randomized distribution; bottom - the real mean difference is lower than most points in the randomized distribution).

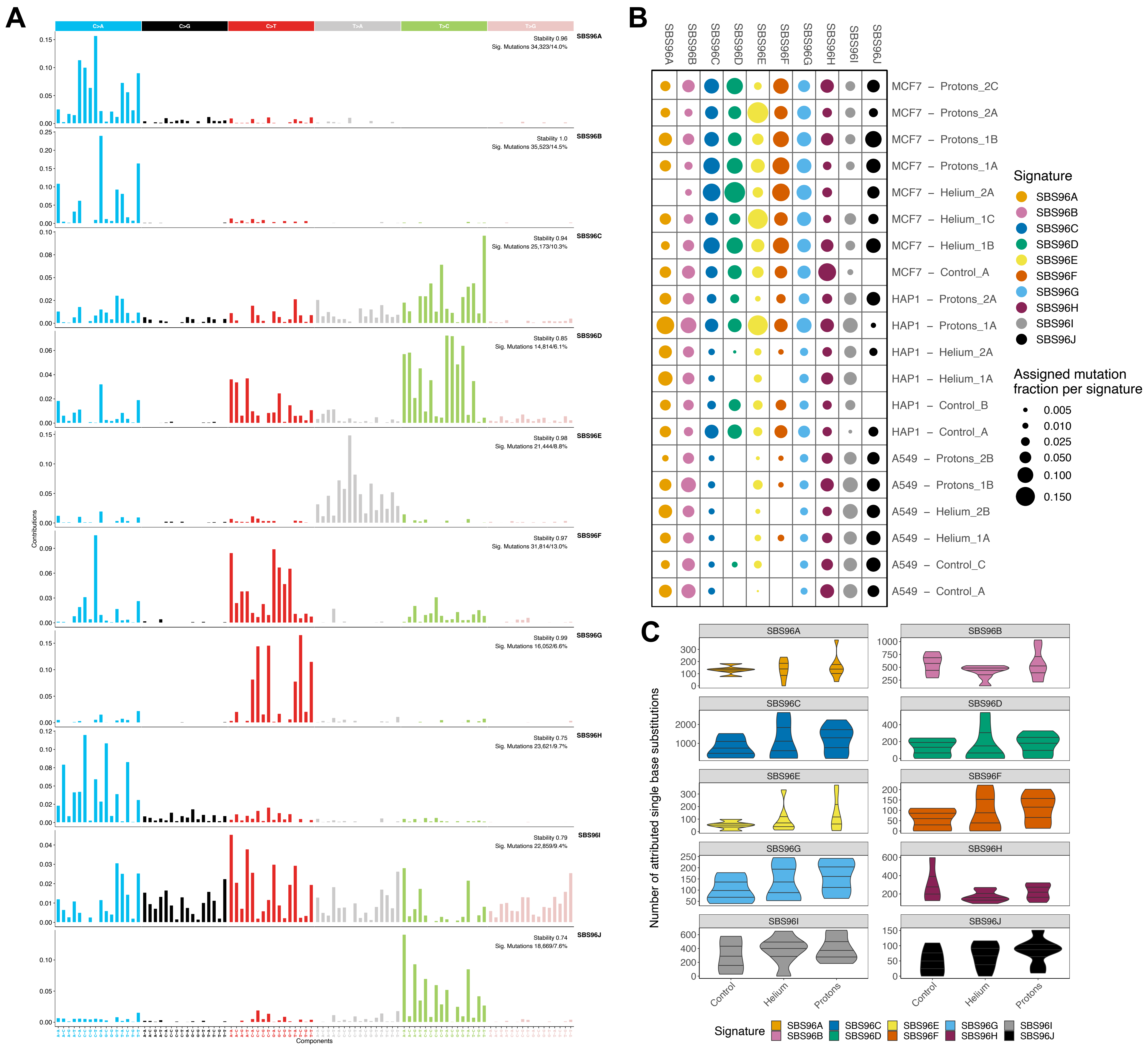

**Supp. Figure 2. Extracted SNV spectra. A.** Mutational spectrum of the 10 SNV signatures extracted using the SigProfiler ExtractorR algorithm. Each signature corresponds to the distribution of point mutations into one of the 96 possible substitution categories taking a trinucleotide context: 6 possible point mutations (taking into account base complementarity, i.e. A=T; G=C) at the target position and 4 at each surrounding genomic location = 6*4*4=96 combinations. **B.** Exposures of all extracted SNV mutational signatures expressed as a fraction of the total mutation burden generated by each signature. **C.** Number of point mutations attributed to each SNV signature for each treatment, pooled by cell line. Black horizontal lines denote the first (lower), second (median) and third (upper) quartiles.

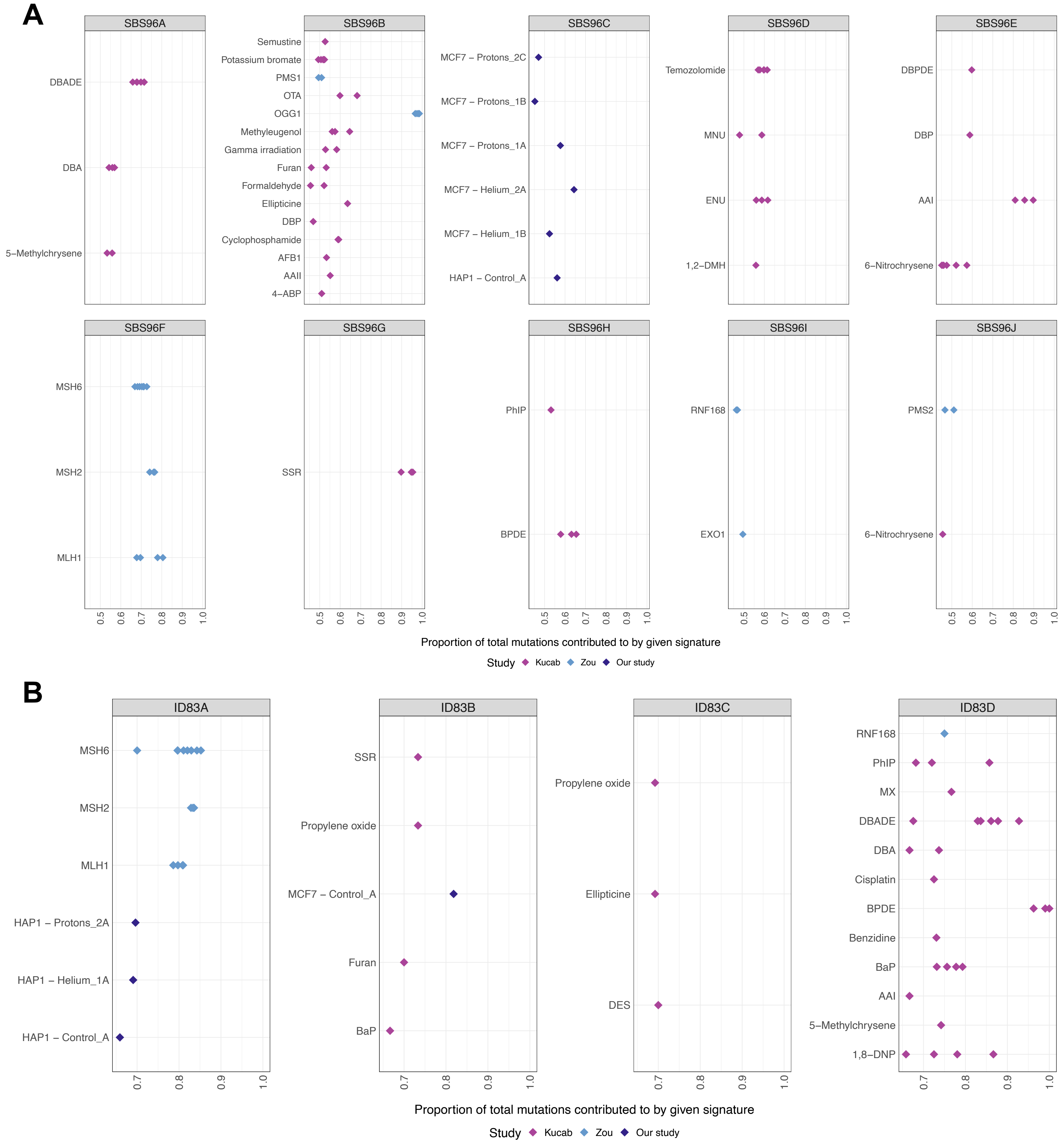


**Supp. Figure 3. High signature exposure in the complete dataset.** Samples from the complete dataset with high exposure to each of the **A.** 10 SNV signatures and **B.** 4 indel signatures. The threshold for high exposure was set at 45% (of total mutations in the sample are generated by the given signature) for SNVs and 65% for indels. The samples were coloured according to their corresponding study: pink for Kucab *et al.* (n=155), blue for Zou *et al.* (n=38), and purple for our cohort (n=21).

A

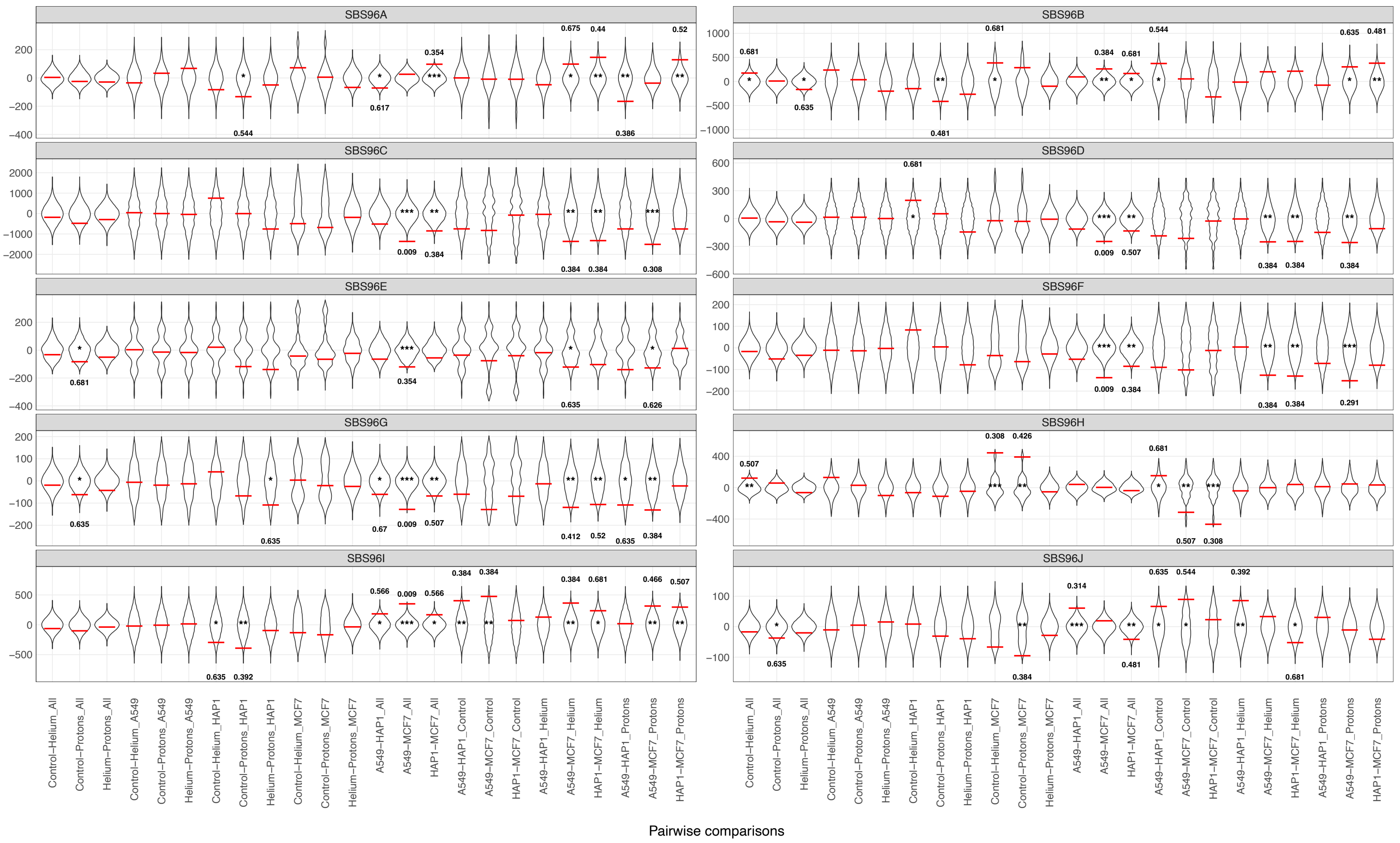


B

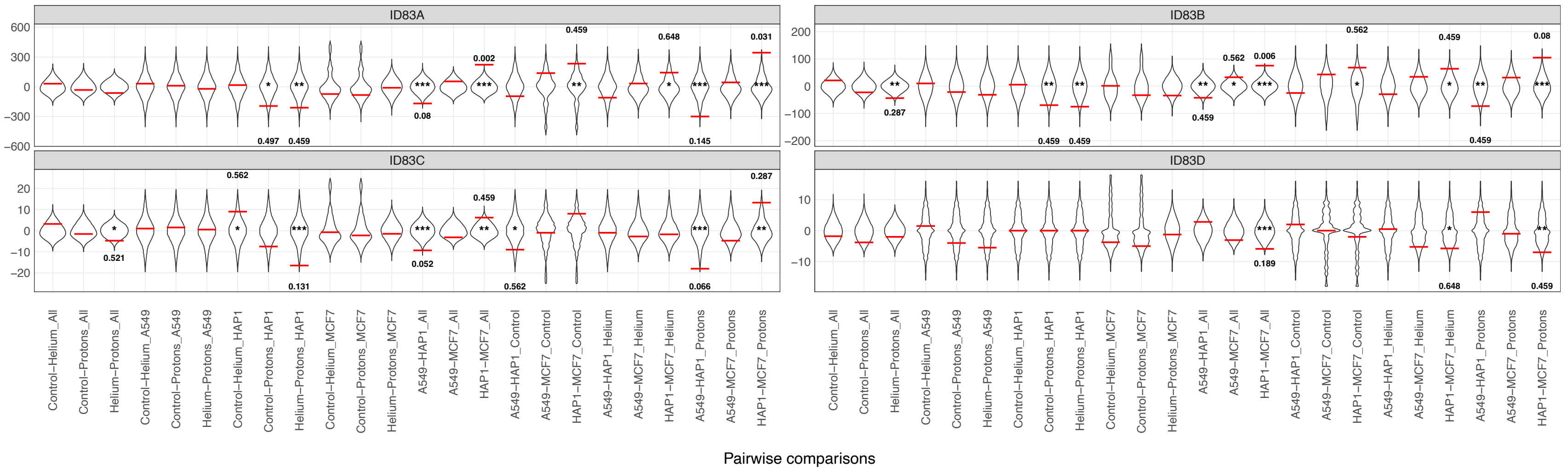


**Supp. Figure 4. Signature exposure randomization tests.** Pairwise differences in treatment/cell line group means (pooled or stratified by treatment or cell line) of signature exposures expressed as raw mutation counts (**A.** SNV signatures, **B.** indel signatures). Red horizontal lines represent the difference between real group means, whereas the violin distributions represent the differences between randomized group means. Unadjusted empirical p-values are marked using the following scale: (*) p <= 0.1, (**) p <= 0.05, (***) p <= 0.01. Q-values are printed for all comparisons with a p-value <= 0.1 on the side of the distribution where the real difference in mean is located.

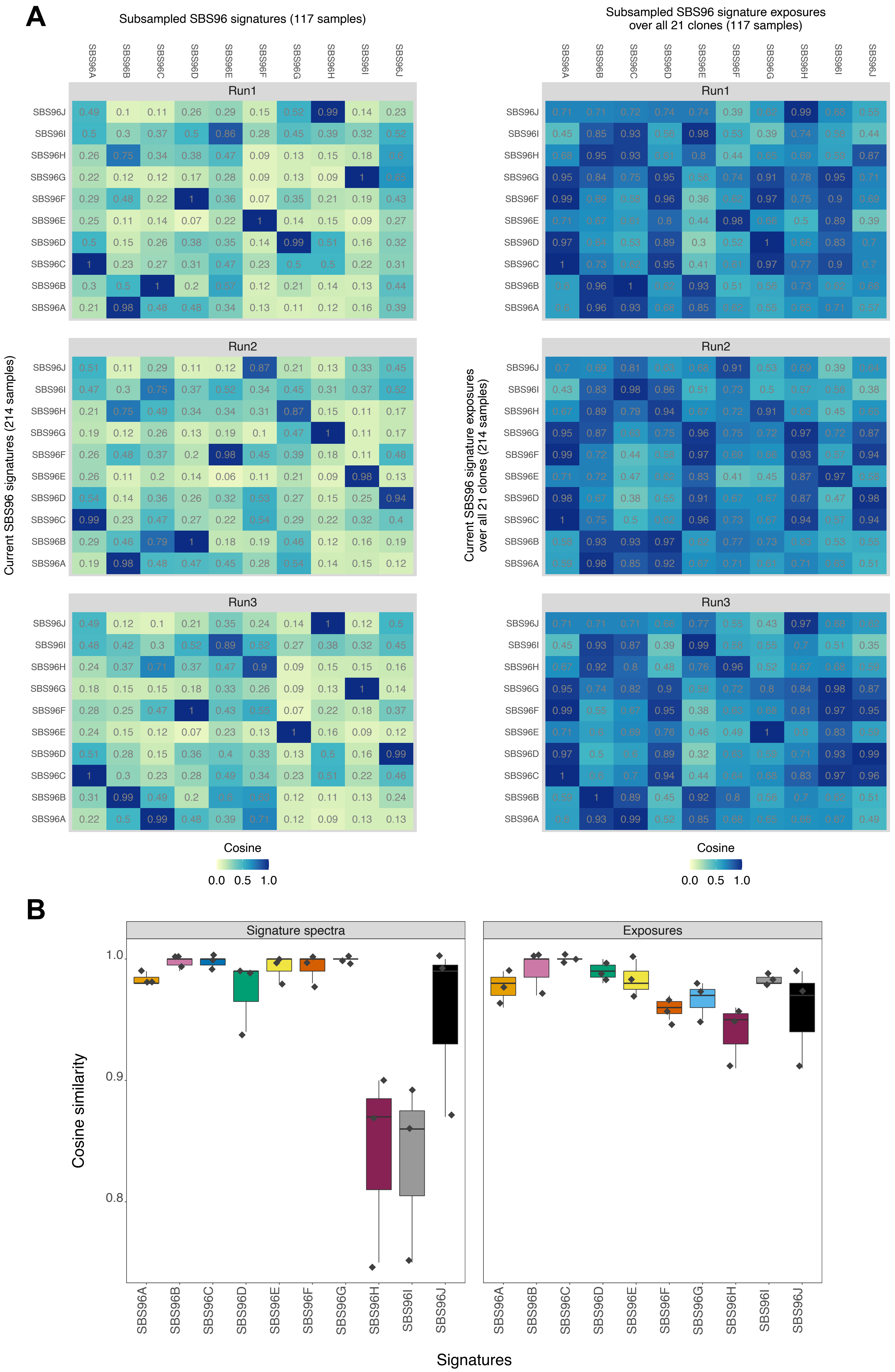


**Supp. Figure 5. Subsampled SNV mutational signatures and exposures. A.** (LEFT) Cosine similarity between the SNV signatures derived from the whole point mutation dataset (n=214) and the spectra derived from three independent subsampling runs (n=117). (RIGHT) Cosine similarity between the SNV signature exposures in our cohort (n=21) derived from the whole point mutation dataset and the exposures from three independent subsampling runs. **B.** (LEFT) Highest signature spectrum cosine similarity values from each subsampling run for each SNV signature in the original set. (RIGHT) Signature exposure (in our cohort) cosine similarity values from each subsampling run (given best signature spectrum match) for each SNV signature exposure in the original set.

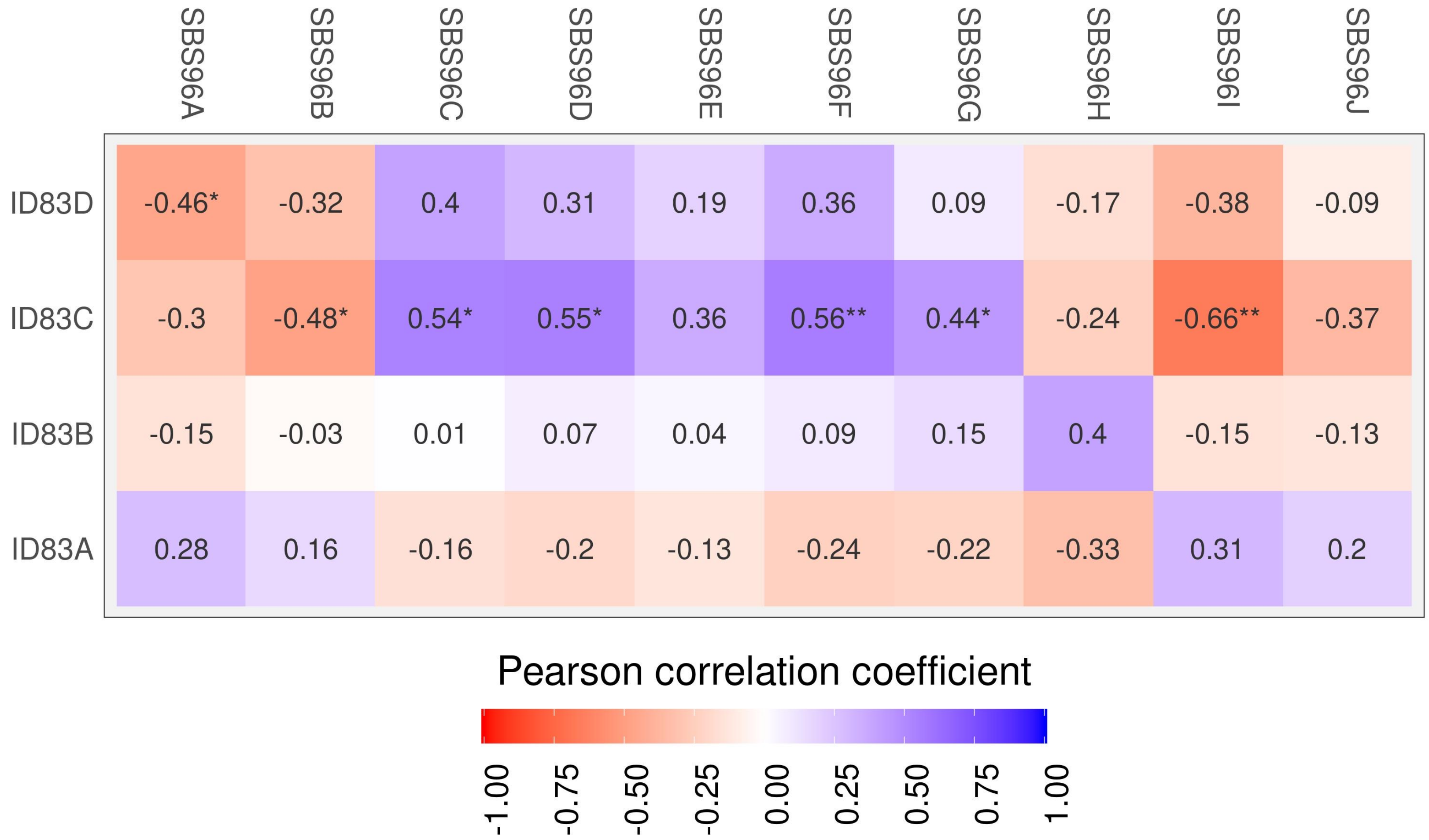


**Supp. Figure 6. Correlation between SNV and indel signature activities.** In each sample we computed, for each SNV signature and indel signature, the proportion of mutations attributed to that particular signature. The plot shows the Pearson correlation between each pair of indel and SNV signature activities in the 21 clones in our cohort. A large positive correlation coefficient (dark blue) suggests that the two signatures in the pair are active in the same samples, whereas a large negative coefficient (dark red) indicates that the activities of the two signatures are inversely correlated (when one signature has high activity the other has low activity and vice versa). A low coefficient (positive or negative) suggests that there is no relationship between the activities of the two signatures in the pair. P-values are marked using the following scale: (*) $p < 0.05$, (**) $p < 0.01$, (***) $p < 0.001$.

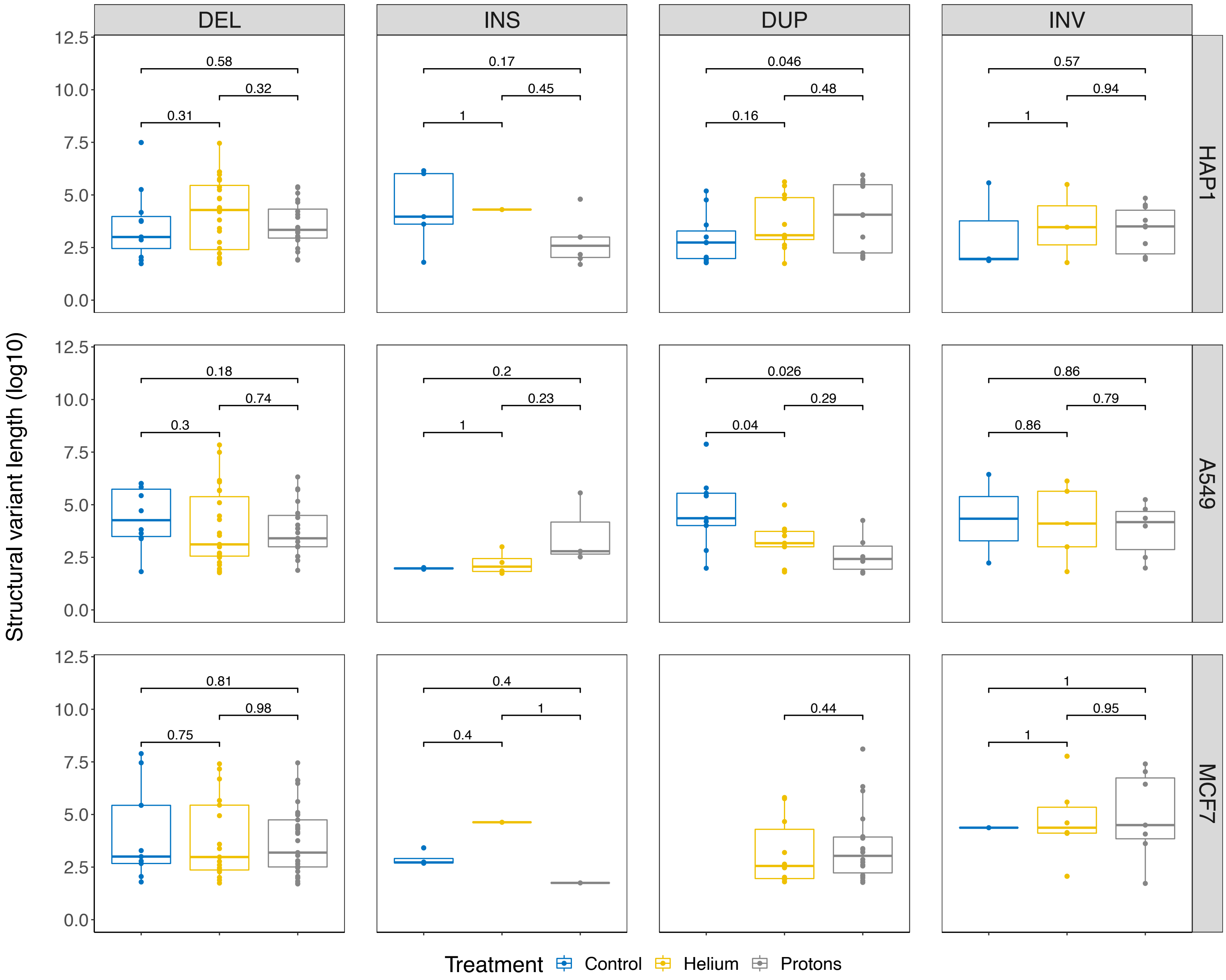


**Supp. Figure 7. Structural variation length distribution.** Distributions of the log10-transformed lengths of structural variants, classified into the 4 main categories, i.e., large deletions, large insertions, duplications and inversions. Rows represented the three different cell lines, A549, HAP1 and MCF7. P-values shown in the plots were computed using the ggpubr R package (t-test mean comparison).

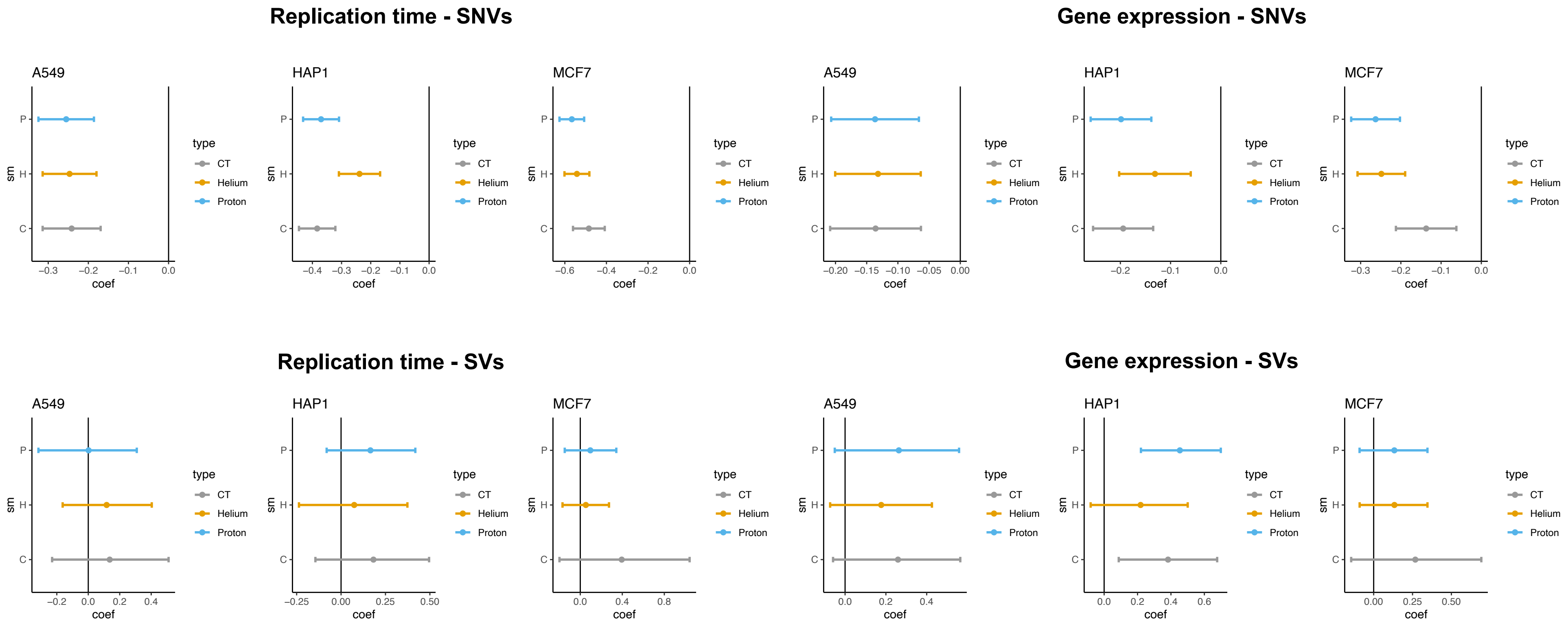


**Supp. Figure 8. Regional mutation rates analysis, comparing negative binomial regression coefficient distribution.** A negative binomial regression was computed for each cell line/condition/clone, in order to detect association between number of mutations (mutation rate) and genomic regions, corrected by trinucleotide context in the case of SNVs. A similar approach than previously developed[55] was applied. The regression was applied to 2 different types of genomic components: replication time and gene expression for SNVs and SVs independently. See Methods for details on regional bins computations.

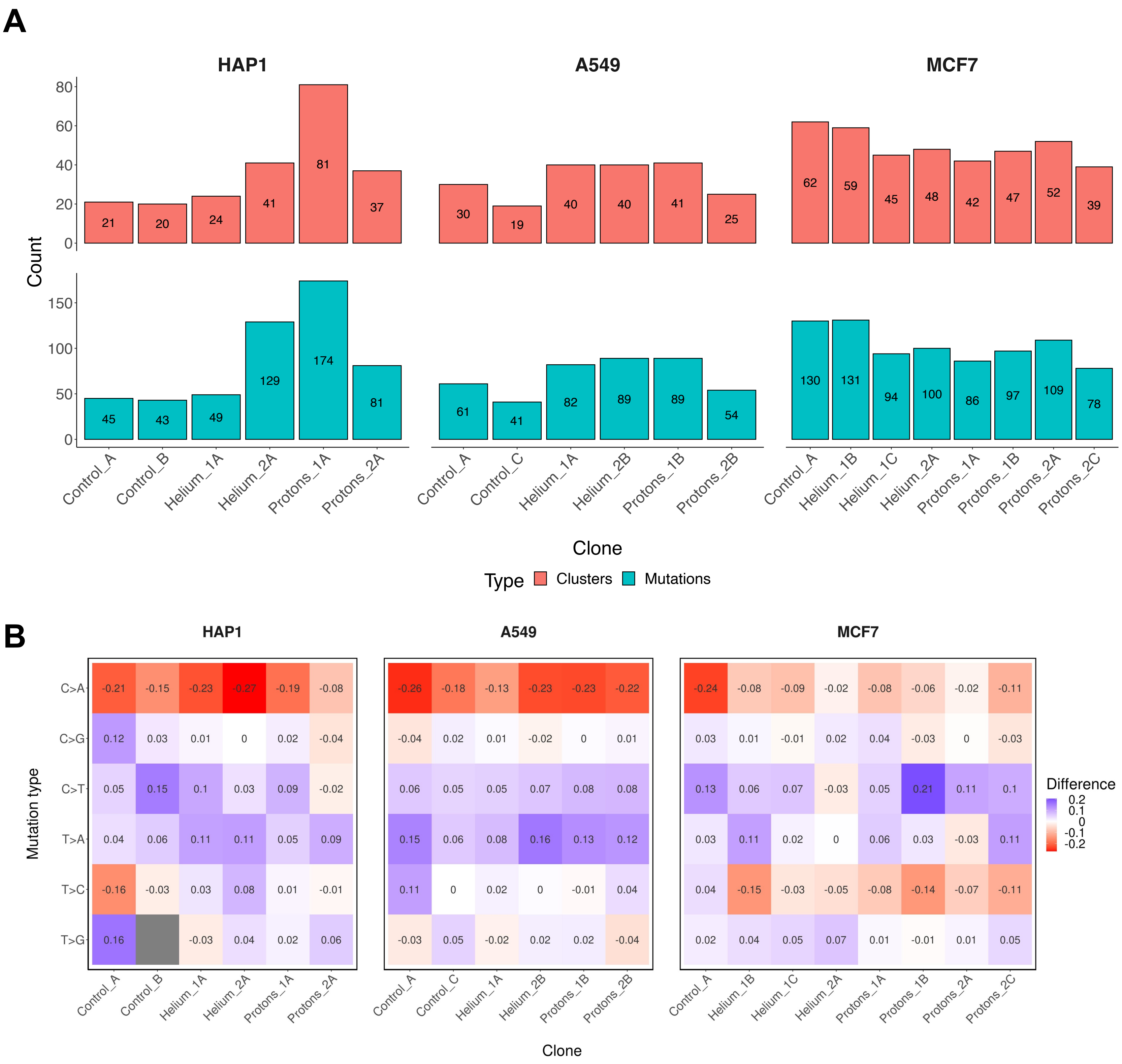


**Supp. Figure 9. Difference in mutational spectrum between mutation clusters and total mutations. A.** Number of total clustered mutations and number of clusters per clone in our cohort. **B.** The relative contribution of each mutation type per clone was calculated for clustered and total SNVs; then, the difference between the clustered and the total contribution was computed. For a large positive value (blue), the mutation type is more abundant in clustered mutations compared to the total, whereas, for a large negative value (red), the mutation type is depleted in clusters compared to the total.
