## Supplementary figures and images for "Proton and alpha radiation-induced mutational profiles in human cells"

### Supplementary Figure 10

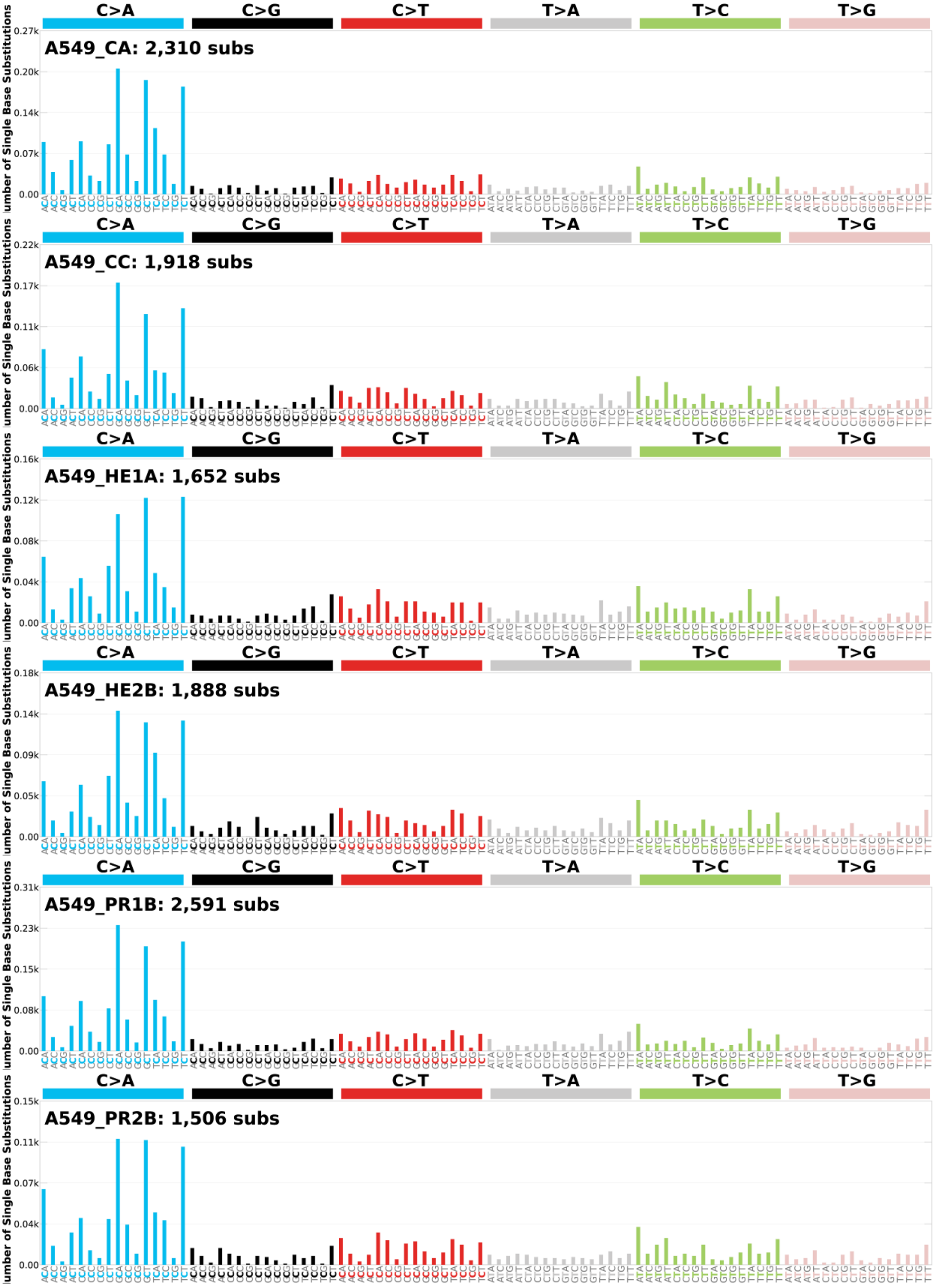

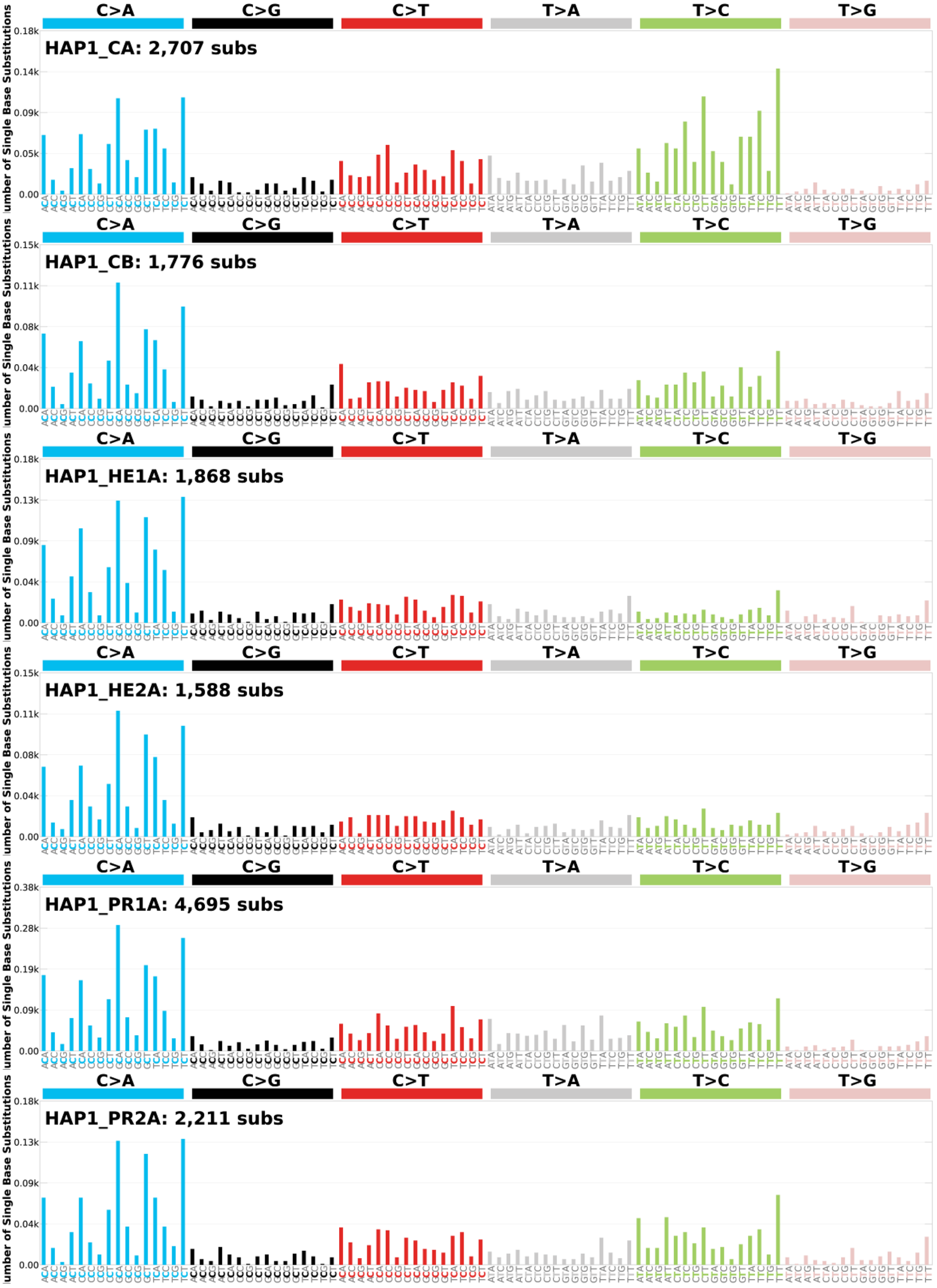

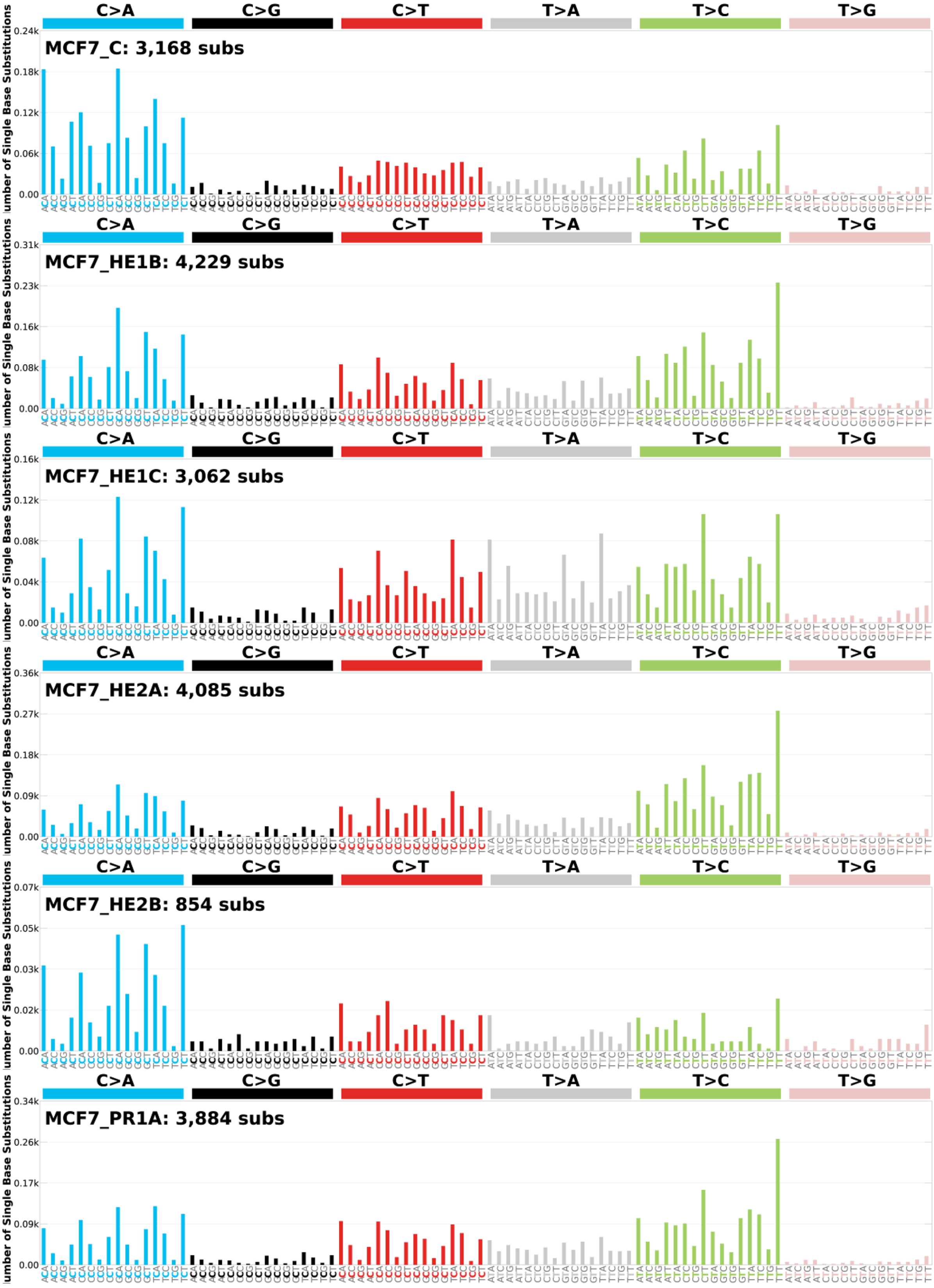

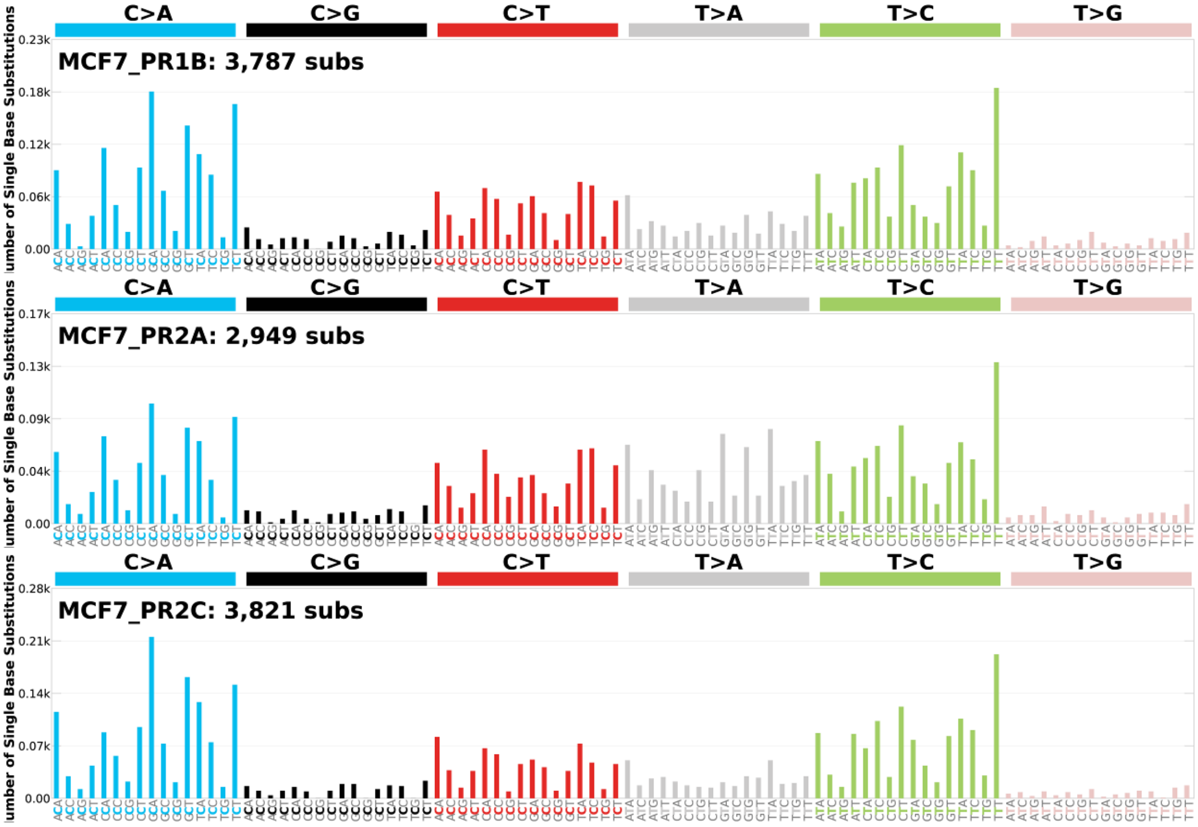


**Supplementary Figure 10.** 96-class spectrum of point mutations in each clone of our cohort.

### Supplementary Figure 11

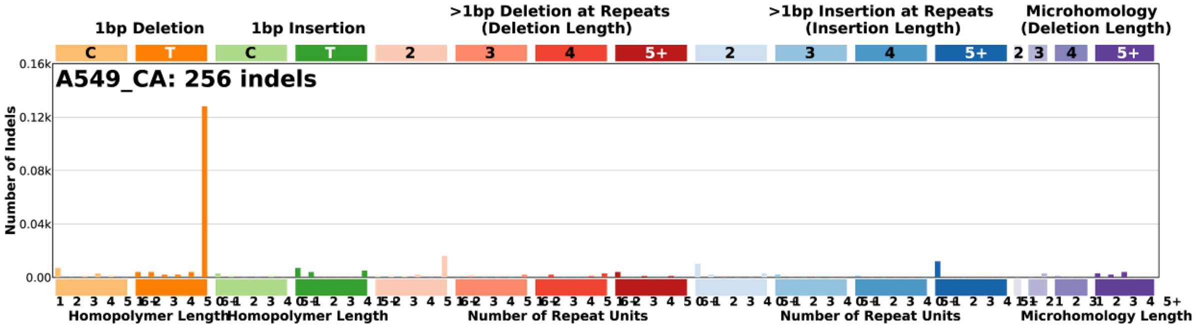

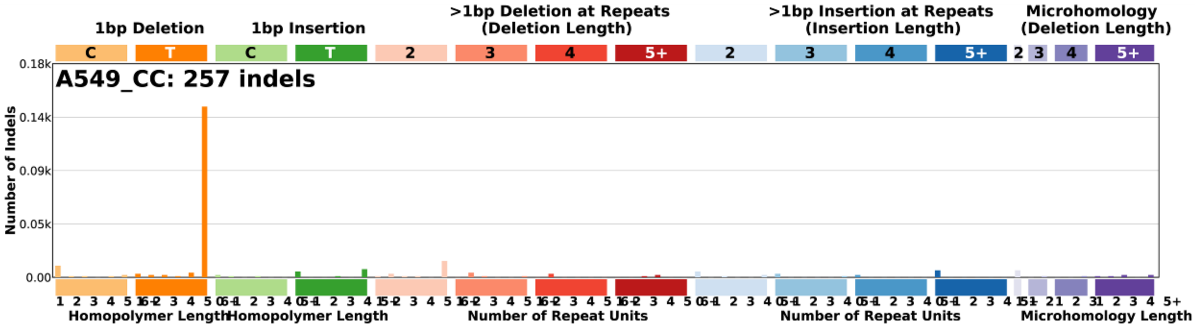

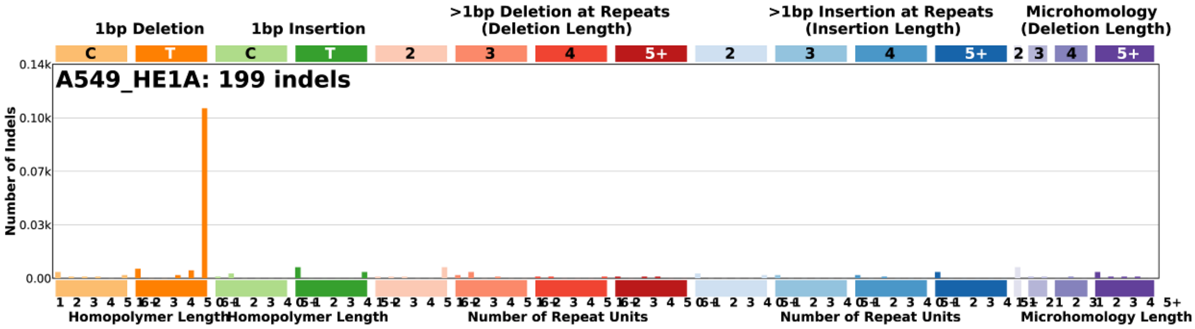

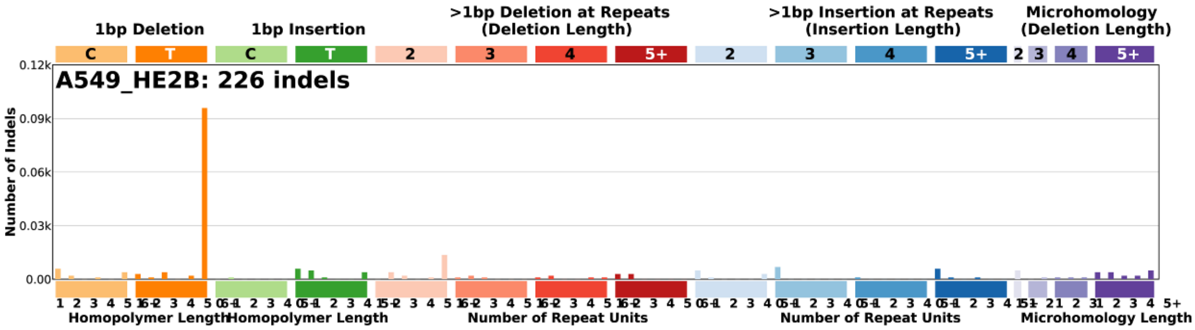

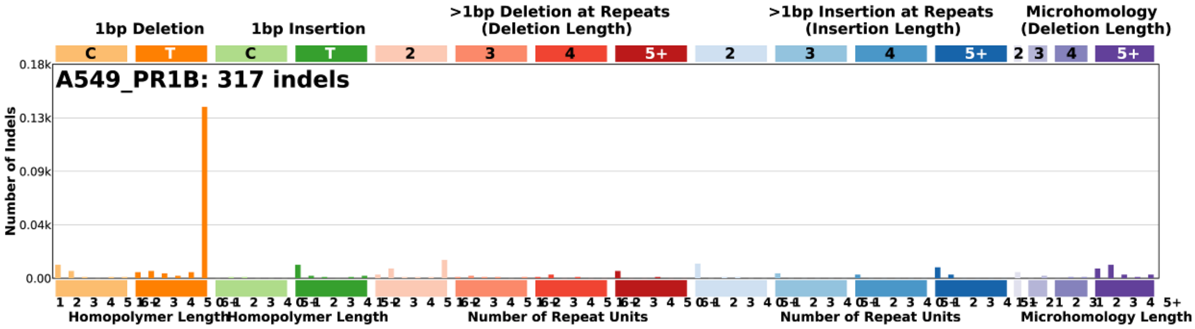

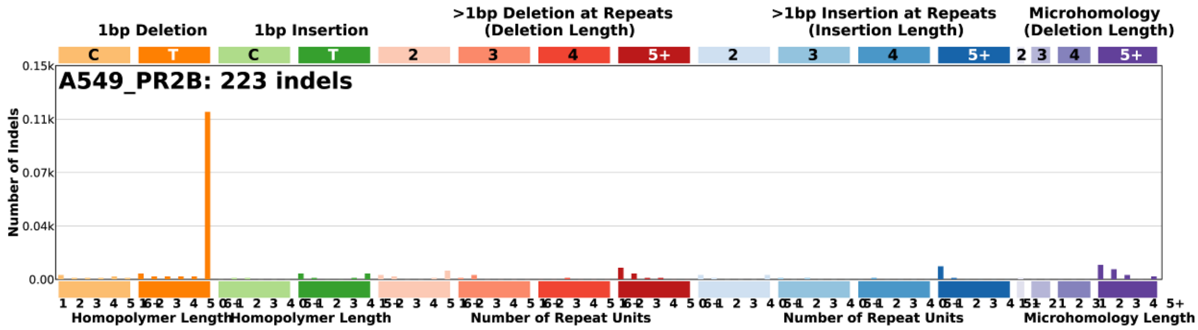

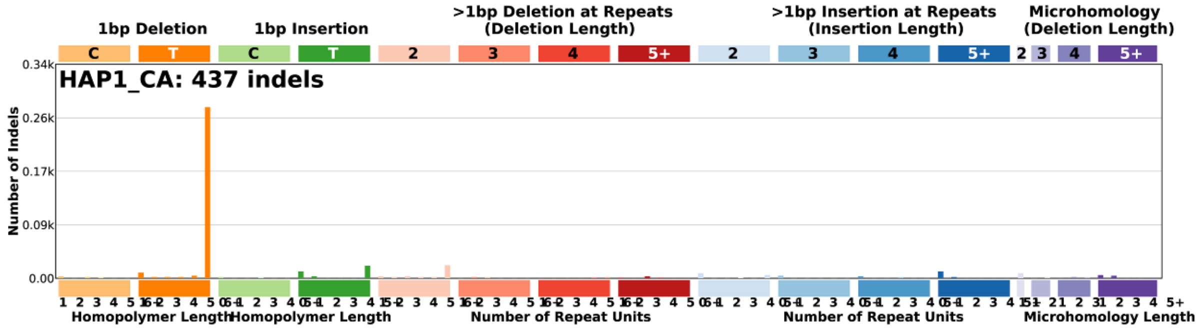

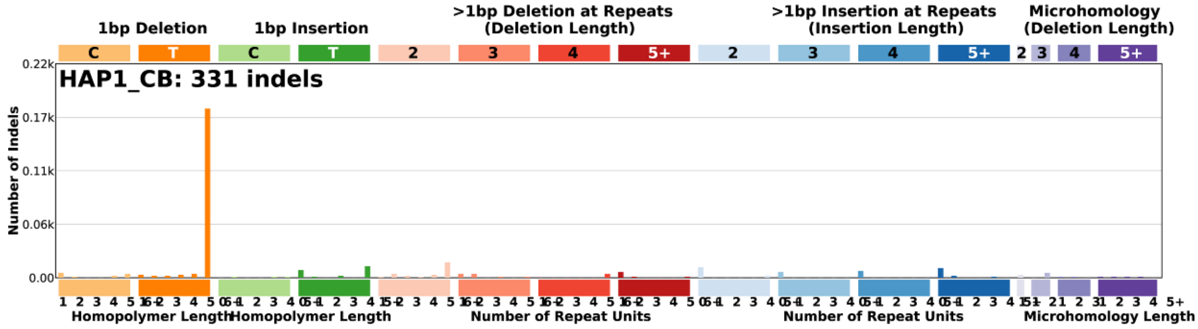

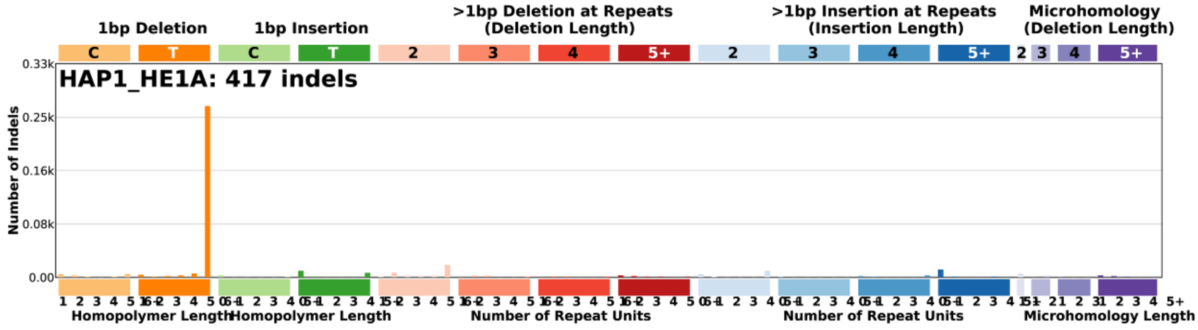

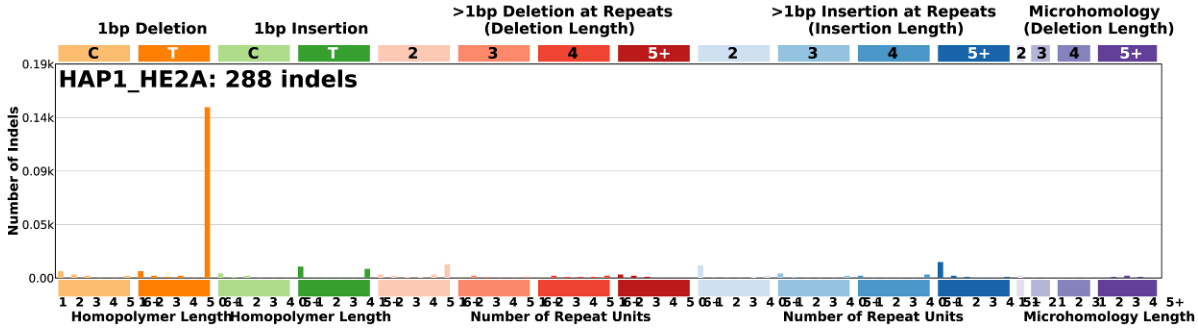

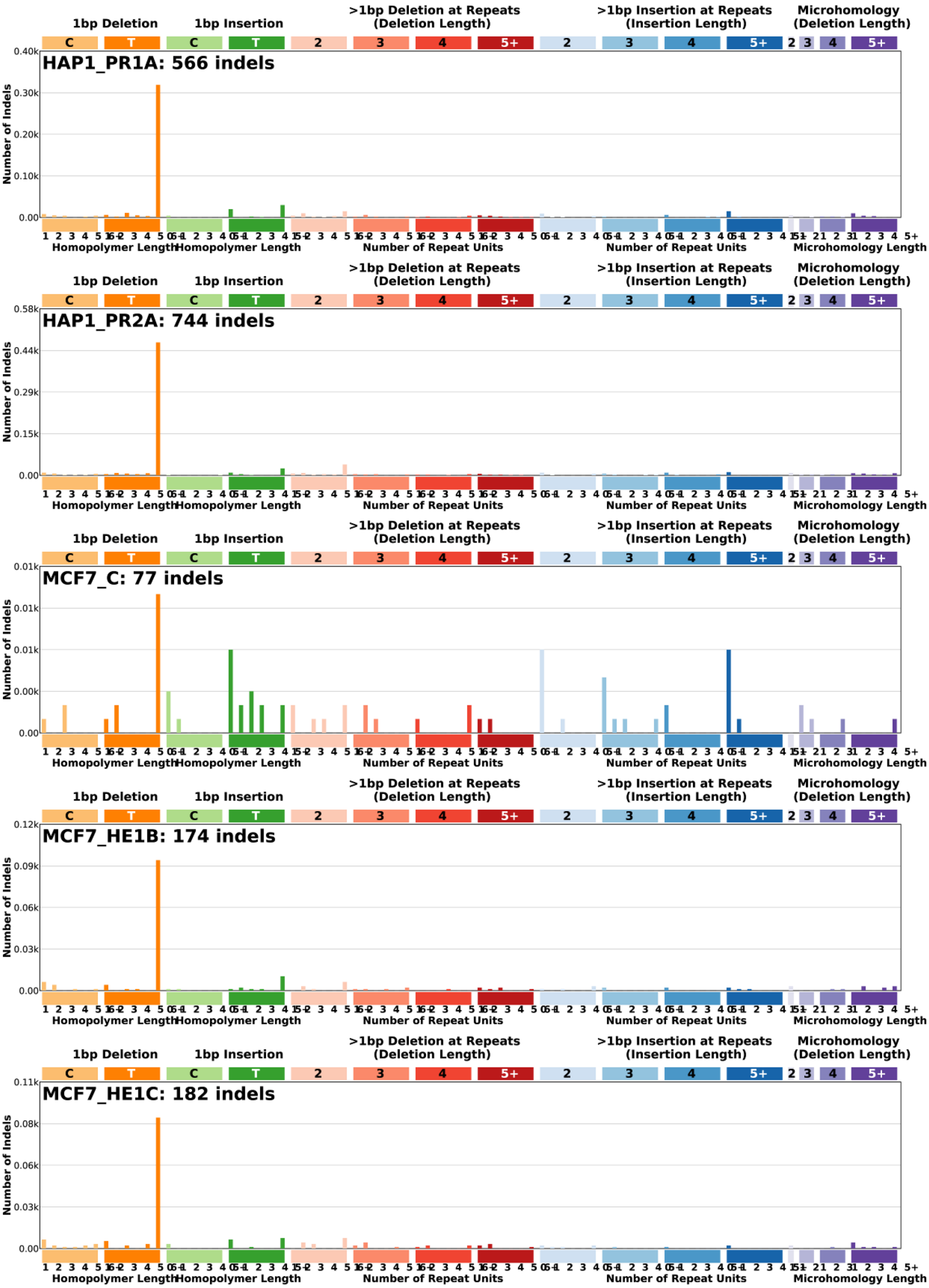

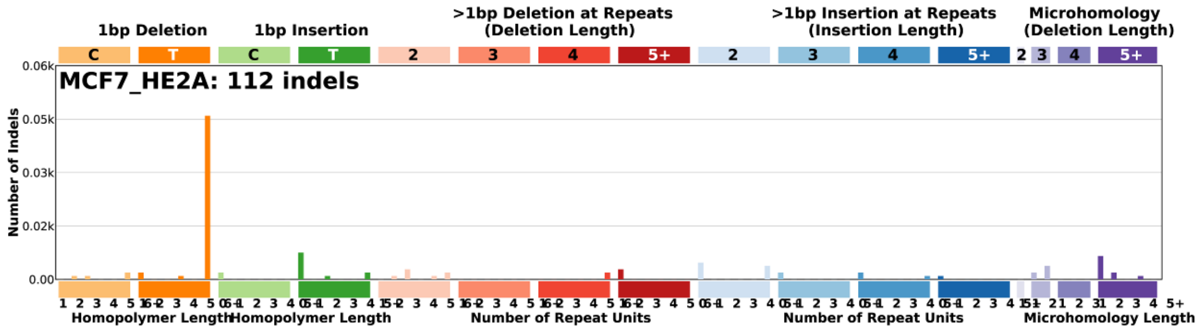

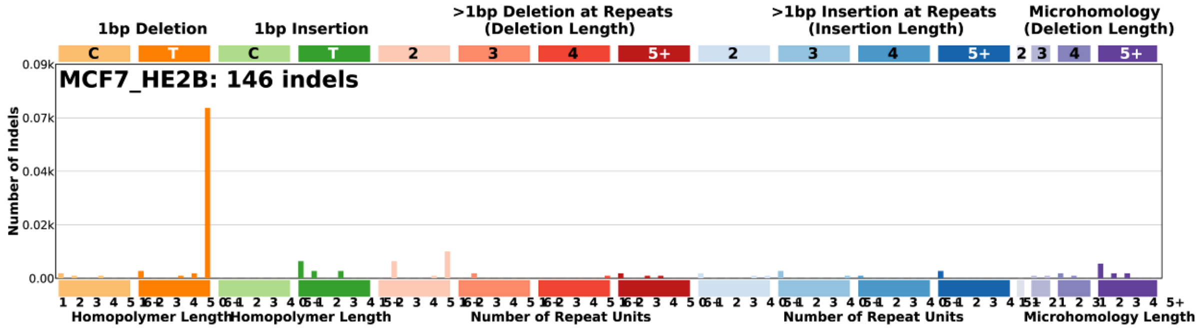

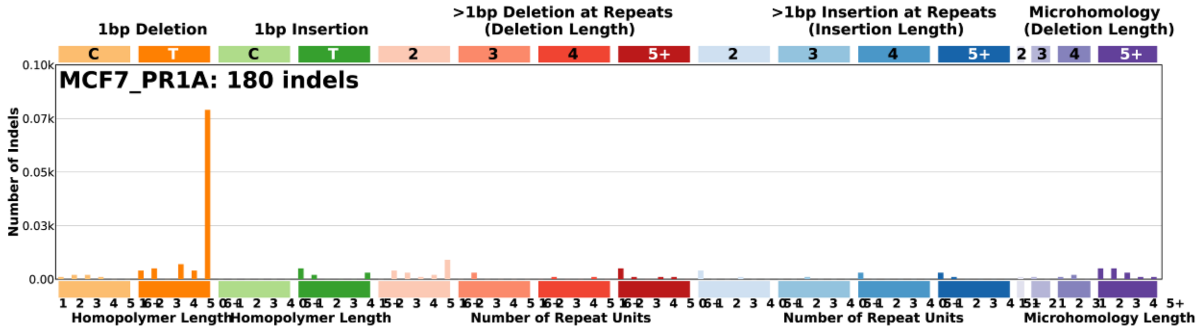

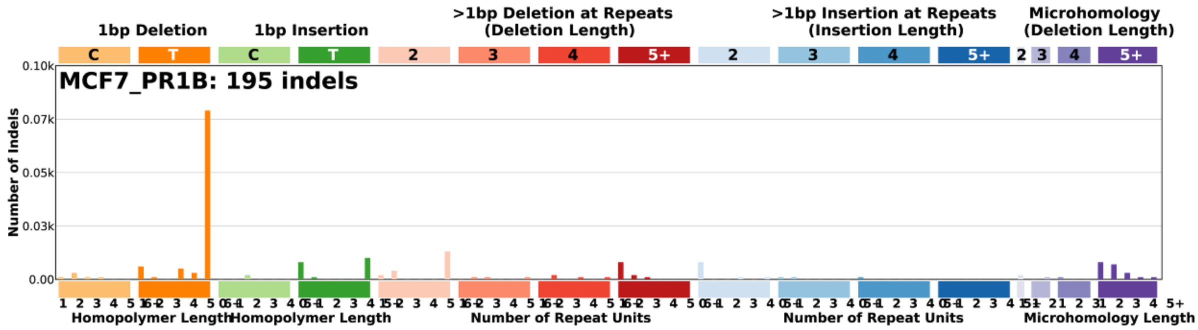

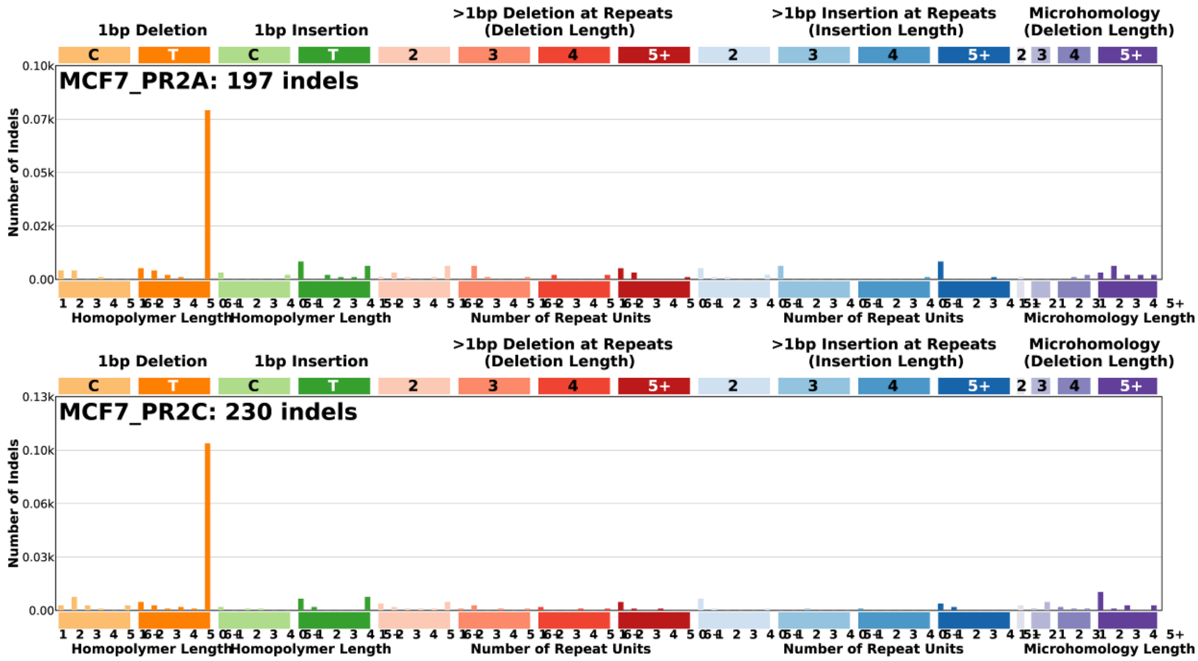


**Supplementary Figure 11.** 83-class spectrum of indel mutations in each clone of our cohort.

### Supplementary Figure 12

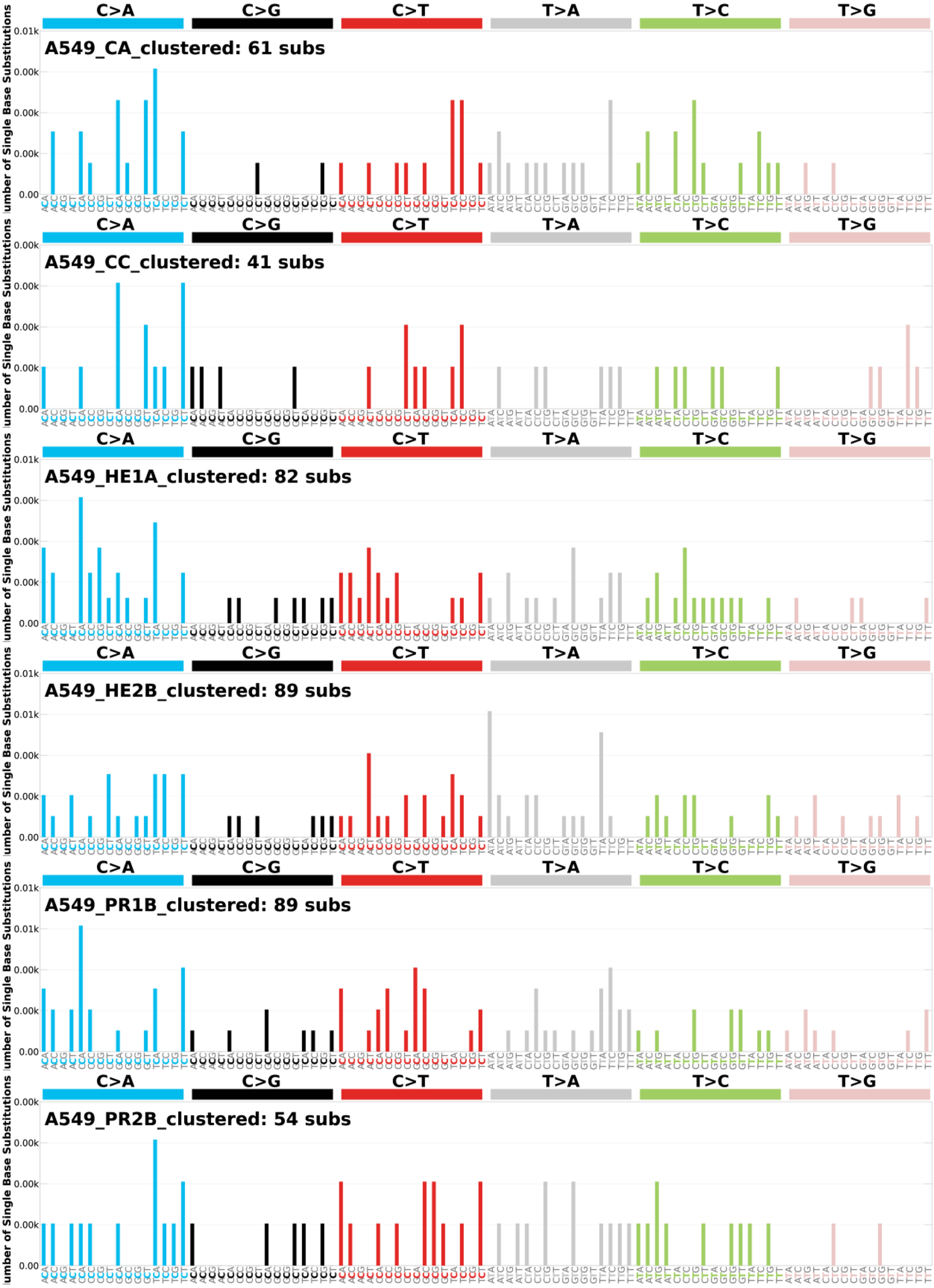

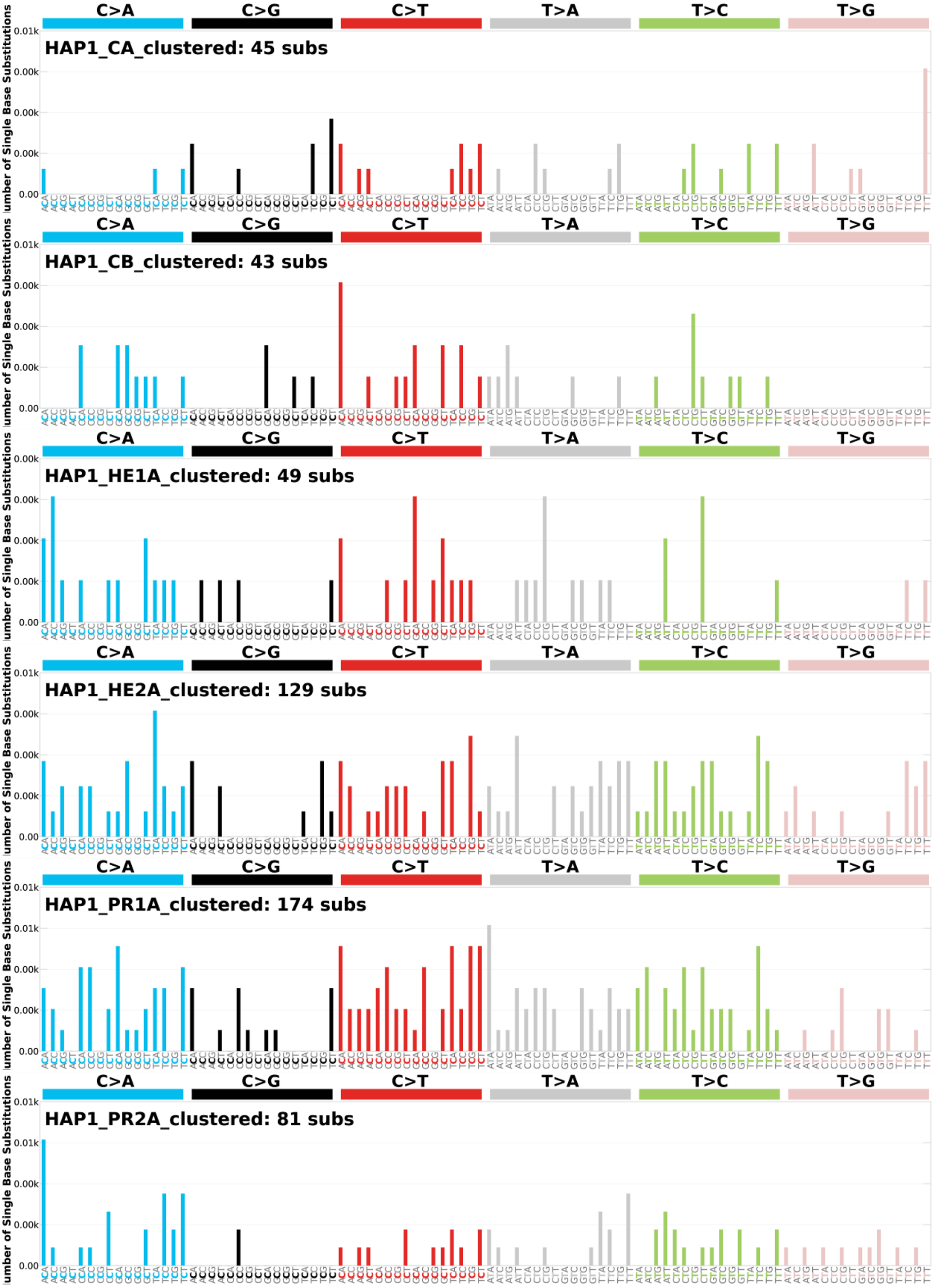

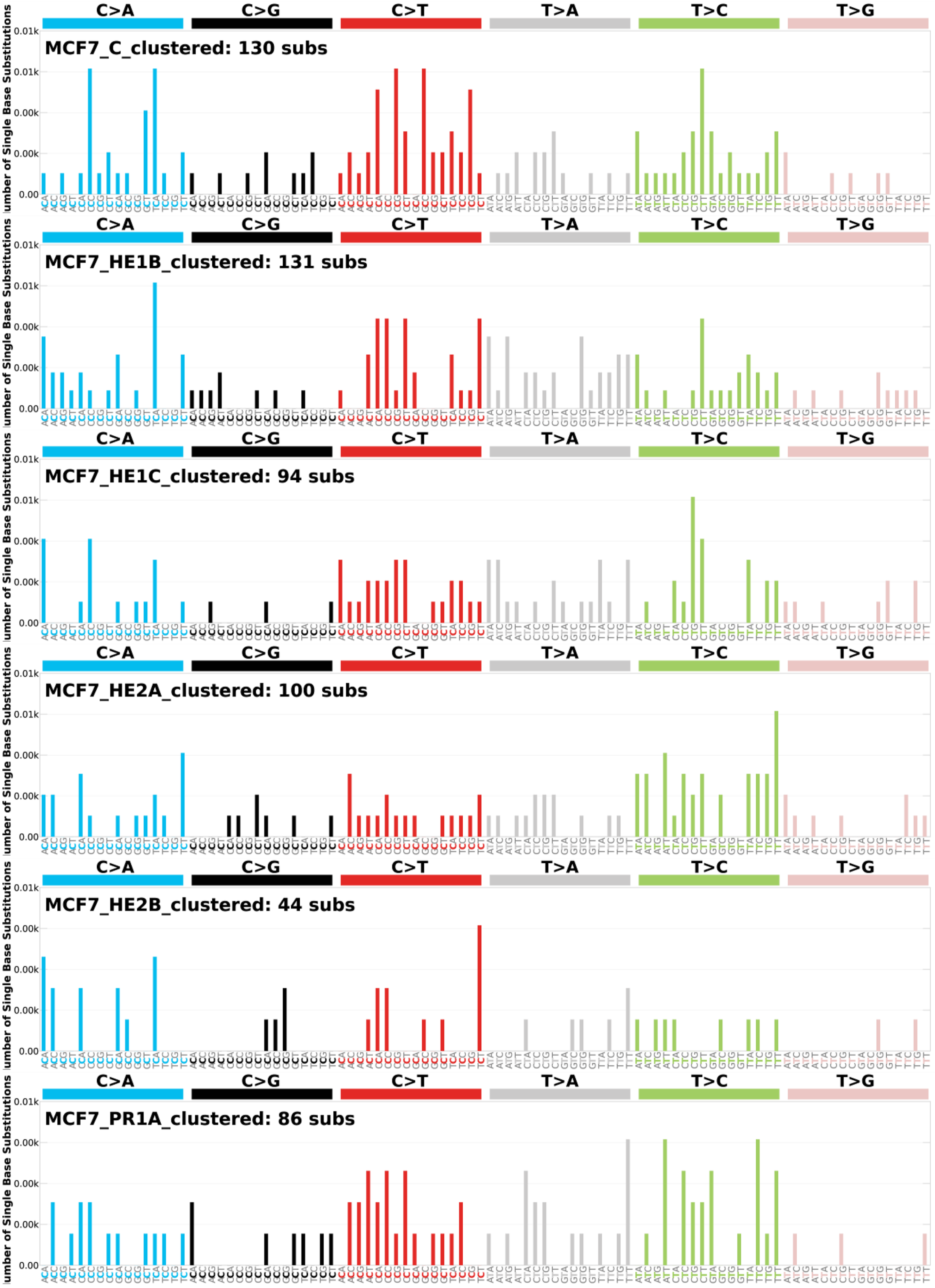

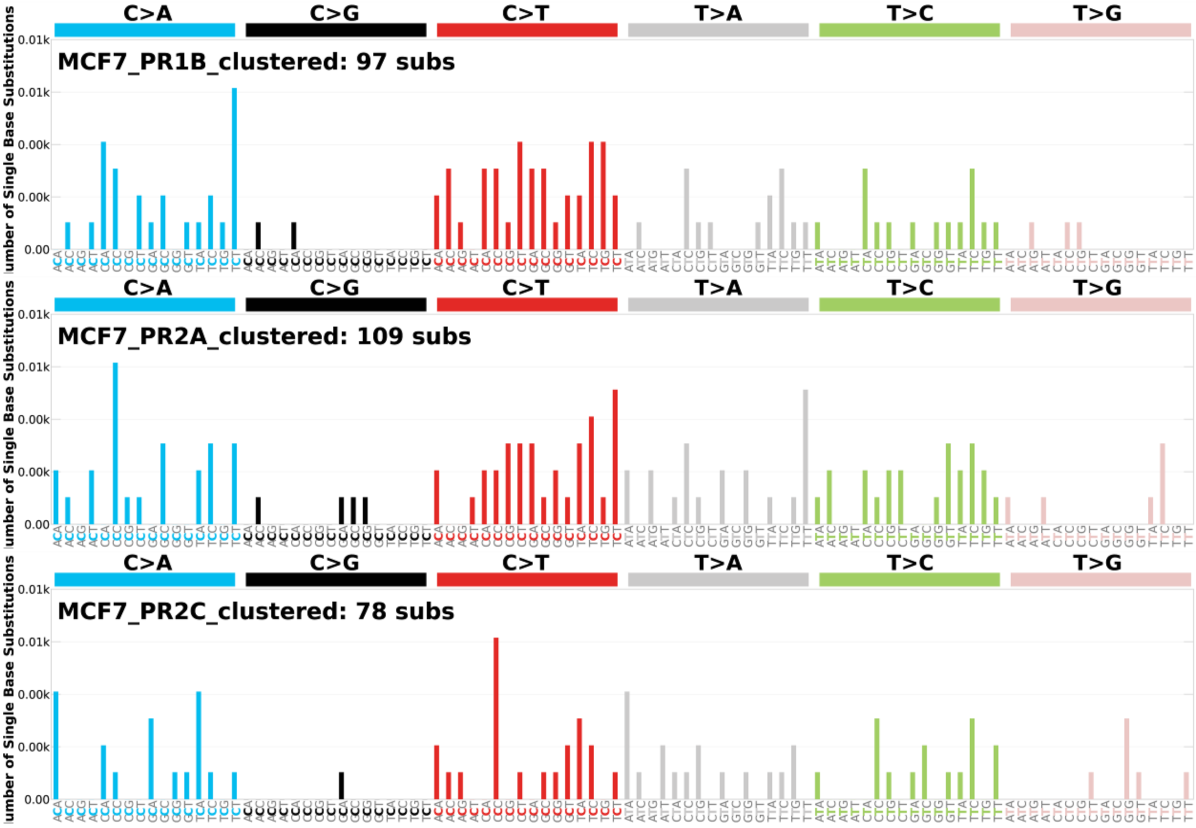


**Supplementary Figure 12.** 96-class spectrum of clustered point mutations in each clone of our cohort.
